# NF-κB-repressed Sirt3 mediates testicular cholesterol metabolism and cytoskeleton assembly via P450scc/SOD2 deacetylation during spermatogenesis

**DOI:** 10.1101/2021.02.22.432399

**Authors:** Mei Wang, Ling Zeng, Yao Xiong, Xiao-fei Wang, Lin Cheng, Fang Wang, Ping Su, Yuan-zhen Zhang

## Abstract

Testicular homeostasis requires the balanced interplay between specific molecules in Sertoli cells, Leydig cells, germ cells. Loss of this coordination can lead to the disruption of spermatogenesis, even male infertility. By operating the upregulation and downregulation of Sirt3 in our male subfertility rats model and two testicular cells models, we indicated that Sirt3 overexpression and activator ameliorated cholesterol metabolism via P450scc deacetylation in Leydig cells, and cytoskeleton assembly via PDLIM1 with SOD2 deacetylation in Sertoli cells and elongating spermatids. In terms of the upstream regulator of Sirt3, the phosphorylation of NF-κB p65^Ser536^ stimulated the nuclear translocation of NF-κB subunits (p50, p65, RelB), which bound to TFBS1 and TFBS2 synchronously in the promoter of Sirt3, repressing Sirt3 transcription. This study demonstrates that NF-κB-repressed SIRT3 acts directly on cholesterol metabolism of Leydig cells and cytoskeleton assembly of Sertoli cells via P450scc/SOD2 deacetylation to regulate sperm differentiation, influencing spermatogenesis, even male fertility.

**Research organism: Rat, mouse**

## Introduction

Spermatogenesis is a well-defined dynamic process that consists of spermatogonial mitosis, spermatocytic meiosis and spermiogenesis (Griswold, 2016). During this highly specialized process, Sertoli cells provide structural and energy support for germ cells differentiation, growth and spermiation (Yokonishi et al., 2020); Leydig cells transform cholesterol to produce testosterone (Li et al., 2016), which is required for meiosis (Deng et al., 2016), releasing mature sperm (O’Donnell et al., 2009), maintaining blood–testis barrier (BTB) (Mruk and Cheng, 2015). Ectoplasmic specialization (ES), as an actin microfilament-rich anchoring device, comprises Sertoli-elongating spermatid interface (apical ES) and Sertoli-Sertoli cell interface (basal ES) (Mruk and Cheng, 2015). The apical ES interacts with the acrosome, reshaping the sperm head, facilitating spermatid morphology and spermiation; the basal ES with the tight/gap junctions constitutes BTB, protecting from external stimulus (Cheng and Mruk, 2012). Except for ES, the microtubule is also a vital cytoskeleton component. In fact, the function of Sertoli cells in nourishing the developing spermatozoa depends on Leydig cells-secreted testosterone (Makela et al., 2019), providing a communication network in testis. Thus, loss of this coordination can lead to the disruption of spermatogenesis, even male infertility.

Sirtuin 3 (SIRT3) is the primary mitochondrial acetyl-lysine deacetylase that modulates various proteins for mitochondrial function, ROS generation, cell death (Pellegrini et al., 2012) and metabolism (Hor et al., 2020; Palomer et al., 2020). SIRT3 cooperates with SIRT1, targeting PGC-1α in antioxidant defense systems (Chen et al., 2018b). Sirt1 deficiency in mice leads to spermatogenesis derangement via meiotic arrest (Heo et al., 2017) and acrosome biogenesis disruption by autophagy (Liu et al., 2017a). Despite the high expression of SIRT3 in mammalian testis (Yu et al., 2014; Yue et al., 2014), the role of SIRT3 in testis remains ambiguous. Our previous study has evidenced that testicular cell injury induced by cadmium (Cd) in rats is associated with mitochondrial autophagy and oxidative stress (Wang et al., 2020). Then, a question appears: whether manipulating Sirt3 can also regulate testicular injury induced by Cd? Even, what’s the functional role of Sirt3 in testis?

Herein, our male subfertility model indicated that SIRT3 activator melatonin (Mel) rescued cholesterol metabolism (and testosterone biosynthesis) by Abca1, Hsd17b3, Scap, Srebf2, Star and F-actin-containing cytoskeleton assembly by ectoplasmic specialization and microtubule-based manchette during spermiogenesis, facilitating sperm motility and count in Cd-treated rat testis. The expression of testosterone biosynthesis-associated markers (NF-κB p65 and P450scc) and cytoskeleton markers (PDLIM1, SOD2), especially SIRT3 presented responsive changes. Further, generating Sirt3 knockdown/overexpression models in TM3 (mouse Leydig) cells and TM4 (mouse Sertoli) cells respectively, we found that like Mel, Sirt3 overexpression ameliorated cholesterol metabolism (and testosterone biosynthesis) via P450scc deacetylation in Cd-treated TM3 cells and cytoskeleton assembly via PDLIM1 with SOD2 deacetylation in Cd-treated TM4 cells. Sirt3 knockdown-induced cell disruption failed to be salvaged by Mel. In terms of the upstream regulator of SIRT3, phosphorylation of NF-κB p65^Ser536^ stimulated the nuclear translocation of NF-κB subunits (p50, p65, RelB), which bound to TFBS1 and TFBS2 in the promoter of Sirt3 synchronously, repressing Sirt3 transcription. Collectively, this study demonstrates the novel role of SIRT3 in testicular cells: **Sirt3 transcriptional repression by NF-κB orchestrates cholesterol metabolism via P450scc deacetylation in Leydig cells; whereas, in Sertoli cells, SIRT3 regulates cytoskeleton assembly via PDLIM1 with SOD2 deacetylation, facilitating sperm differentiation during spermiogenesis.** Our findings shed light on the multi-faceted action of SIRT3 and establish a novel signaling network in the crosstalk between testicular different cells. Understanding the role of SIRT3 in cholesterol metabolism and cytoskeleton assembly will contribute to the development of spermatogenesis. SIRT3 may serve as a potential therapeutic target for spermatogenesis disorganization, male infertility, even other metabolic diseases.

## Results

### SIRT3 activator Mel rescues male reproductive function injury induced by Cd in adult rats, including cholesterol metabolism and structure disruptions

To investigate whether SIRT3 was involved in the testicular function, we first examined the effects of SIRT3 activator Mel by a previous subfertility model in Cd-treated adult male SD rats. Results showed that Mel significantly reversed sperm count, sperm motility, serum testosterone reduction induced by Cd (**Fig. 1a-c**). The concentration of Cd in the testis was apparently reduced by Mel (**Fig. 1d**). Like other steroid hormones, testosterone was biosynthesized from cholesterol, so we explored the levels of serum total cholesterol (TC) and free cholesterol (FC). Interestingly, Cd provoked the augment of serum TC and FC rather than the decline of those, but Mel overturned the above changes (**Fig. 1e, f**).

**Fig. 1.**
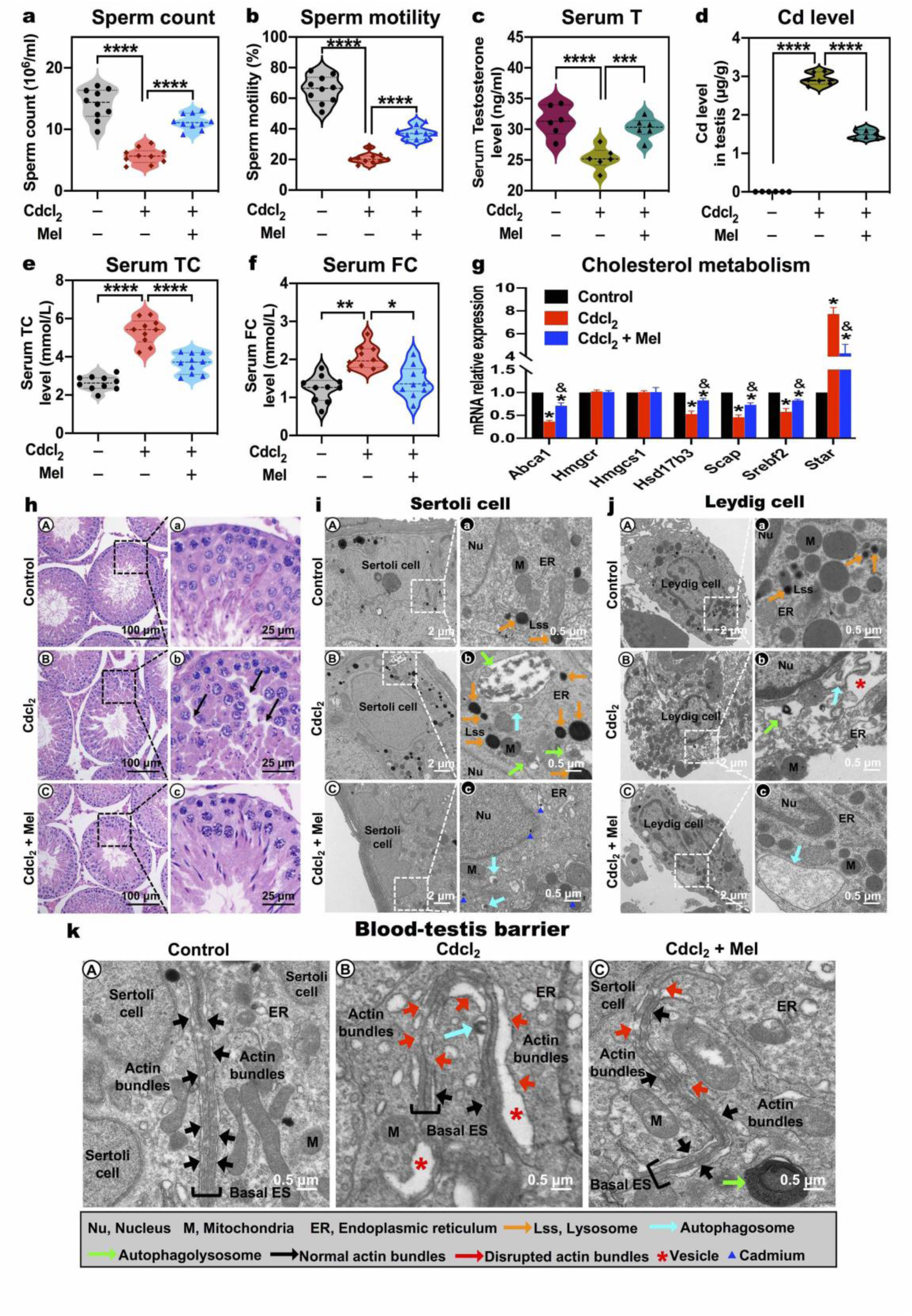
SIRT3 activator Mel rescues male reproductive function injury induced by Cd in adult rats, including cholesterol metabolism and structure disruptions. **a** Sperm count. **b** Sperm motility. **c** Serum testosterone (T) level. **d** Cadmium (Cd) level in testis. **e** Serum total cholesterol (TC) level. **f** Serum free cholesterol (FC) level. **g** mRNA expression levels of cholesterol metabolism markers of Abca1, Hmgcr, Hmgcs1, Hsd17b1, Scap, Srebf2, Star in testis. **h** Histological results of the testis in rats. **i-k** Representative transmission electron micrographs (TEM) depicting the ultrastructure of Sertoli cell, Leydig cell, blood-testis barrier (BTB) in testis. Areas for the cell in the left column (Scale bar, 2 μm) have been shown for further details in the right column (Scale bar, 0.5 μm) in **i** and **j**. Nu, nucleus; **M**, mitochondria; ER, endoplasmic reticulum; orange arrowheads indicate lysosome; turquoise arrowheads indicate autophagosome; spring arrowheads indicate autophagolysosome; black arrowheads indicate normal actin bundles in BTB; red arrowheads indicate disrupted actin bundles in BTB; red asterisk indicates vesicle; blueberry triangles indicate potential Cd in testis. \**p* < 0.05; \*\**p* < 0.01; \*\*\**p* < 0.001; \*\*\*\**p* < 0.0001 in a-f. \**p* < 0.05 compared with the control. &*p* < 0.05 compared with the Cd-treated group in **g**.

To elucidate the reason, we checked cholesterol metabolism markers, including a series of steroidogenic enzymes, which are responsible for testosterone biosynthesis. Cd suppressed the expression of cholesterol efflux marker Abca1, which was reversed by Mel (**Fig. 1g**), suggesting that Mel could regulate cholesterol efflux blocked by Cd in testicular cells. Cholesterol biosynthesis markers Hmgcr and Hmgcs1 (Luo et al., 2020) didn’t present any differences in Cd with or without Mel group when compared to the control group (**Fig. 1g**), indicating that whether Cd or Mel was independently associated with cholesterol biosynthesis in the testis. Besides, the expression of Star, a key transporter of cholesterol from cytoplasm into the mitochondria for testosterone biosynthesis (Rubinow, 2018), was dramatically increased in Cd-treated group, and Mel reversed the augment of Star in the testis (**Fig. 1g**). As predicted, more cholesterol could be delivered into the mitochondria, and then produced more testosterone in Cd-treated group. Nonetheless, the fact was that Cd diminished testosterone production. Given that Scap/SREBP system was sensitive to the concentration of cholesterol in the plasma membrane and regulated genes transcription of cholesterol metabolism (Das et al., 2014), we subsequently inspected the expression of Scap and Srebf2, which were significantly dwindled in Cd-treated group and raised back in Cd with Mel group (**Fig. 1g**). Hsd17b3, as an indispensable enzyme for testosterone biosynthesis, specifically catalyzed the conversion of androstenedione to testosterone (Marshall et al., 2002). Mel salvaged the reduction of Hsd17b3 expression induced by Cd (**Fig. 1g**). Above results revealed that SIRT3 activator Mel rescued cholesterol metabolism disruption in male subfertility model induced by Cd.

Mel decreased the morphologic damage of the seminiferous tubules was decreased by Mel. As depicted in **Fig. 1h**, testicular cells in Cd-treated group were arranged loosely with a large number of voids (**Fig. 1h (Bb)**), while whether the control group or Cd with Mel group exhibited an intact and dense lumen structure (**Fig. 1h (Aa, Cc)**). For further exploring which of testicular cells were affected in this study, we analyzed the ultrastructure of the testis by transmission electron microscopy (TEM). Both Sertoli cells and Leydig cells presented complete nuclear membrane structure, and normal organelles (such as mitochondria, endoplasmic reticulum) with a low number of lysosomes in the control group (**Fig. 1i (Aa), j (Aa)**). By contrast, Cd apparently induced lots of lysosomes, autophagosomes and enormous autophagolysosomes in Sertoli cells (**Fig. 1i (Bb)**), which were consistent with autophagy observed in previous study (Wang et al., 2020). However, Cd with Mel group showed nearly normal organelles with sporadic autophagosomes, and tiny black unidentified objects (probably Cd) (**Fig. 1i (Cc)**). In Leydig cells, Cd evoked massive cytoplasmic vacuolization, while Mel obviously protected against the abnormal ultrastructure induced by Cd (**Fig. 1j (Bb, Cc)**). Given that Leydig cells were the primary source of testosterone production (Makela et al., 2019), the ultrastructural changes in Leydig cells corresponded to above cholesterol metabolism and testosterone biosynthesis regulation. Our previous study indicated that Cd disturbed BTB via oxidative stress (Chen et al., 2018a), so we asked whether Mel could protect from BTB injury. The basal ES and other junctions constituted the BTB between adjacent Sertoli cells. The control group exhibited adjacent Sertoli cells and normal basal ES with the actin bundles; Cd disrupted the basal ES and actin bundles with large vacuoles (asterisks), but Mel rescued BTB disruption induced by Cd (**Fig. 1k**). Actually, the disruption of BTB led to germ cell loss and male infertility (Cheng and Mruk, 2012), which might account for sperm count reduction.

Above results proved that Mel regulated Sertoli cells, Leydig cells and BTB, intimating that SIRT3 might engage in cholesterol metabolism and testosterone synthesis by Leydig cells, and affect sperm count by BTB and Sertoli cells.

### SIRT3 activator Mel rescues cytoskeleton (apical ES and microtubule-based manchette) assembly disruption induced by Cd in the elongating spermatid during spermiogenesis

Except for sperm count, Mel rescued sperm motility reduction induced by Cd (**Fig. 1b**). The cytoskeleton assembly of apical ES and microtubule-based manchette in sperm head and flagellum during spermiogenesis, is required for sperm motility acquirement (Pleuger et al., 2020). To test the hypothesis if SIRT3 activator Mel protects sperm motility by regulating cytoskeleton assembly, we screened the spermatid deformation process step by step during spermiogenesis through TEM analysis.

Spermiogenesis begins with round spermatid elongation at step 8, so our observation started from this stage. There were no significant differences between Cd with or without Mel group and the control group at step 8 (**Fig. 2a (A-C)**). From step 9 to 10, bundles of microtubules were assembled to form the manchette structure (black arrow), which was tightly arranged in different groups (**Fig. 2a (D2-F2)**). The apical ES (white arrow), an imperative component for shaping sperm head, was constituted by actin bundles surrounding the acrosome’s outer layer. We found that the sperm head was disturbed with large vacuoles (red asterisk) in Cd-treated group, but Mel reversed the disruption of apical ES and acrosome induced by Cd (**Fig. 2a (D1-F1)**). In the early elongating spermatids of step 10-11, a well-assembled sperm head and long microtubule-based manchette could be observed in the control group (**Fig. 2a (G)**); whereas, in the Cd-treated group, apical ES disappeared with large vacuoles in sperm head, and the manchette was undermined with short microtubules (red arrow) (**Fig. 2a (H)**); however, in Cd with Mel group, apical ES and manchette were slightly impaired (**Fig. 2a (I)**). In the late elongating spermatids of steps 14 to 15, well assembled apical ES and acrosome were finely arranged surrounding the condensed nucleus, and the long manchette could be detected clearly (**Fig. 2a (J)**); instead, in Cd-treated group, gigantic vacuoles replaced the well apical ES, only round dots (red arrow) rather than long microtubules were observed (**Fig. 2a (K)**); yet Cd with Mel group presented slightly apical ES and manchette injury (**Fig. 2a (L)**). Besides, flagella axonemes (yellow arrow), which were associated with sperm motility, were identified. We did not discover any differences in axoneme between three groups, only with some autophagosomes around the axoneme in Cd-treated group (**Fig. 2a (M-O)**). Ultrastructural results displayed that Mel could extricate apical ES and manchette injury caused by Cd.

**Fig. 2.**
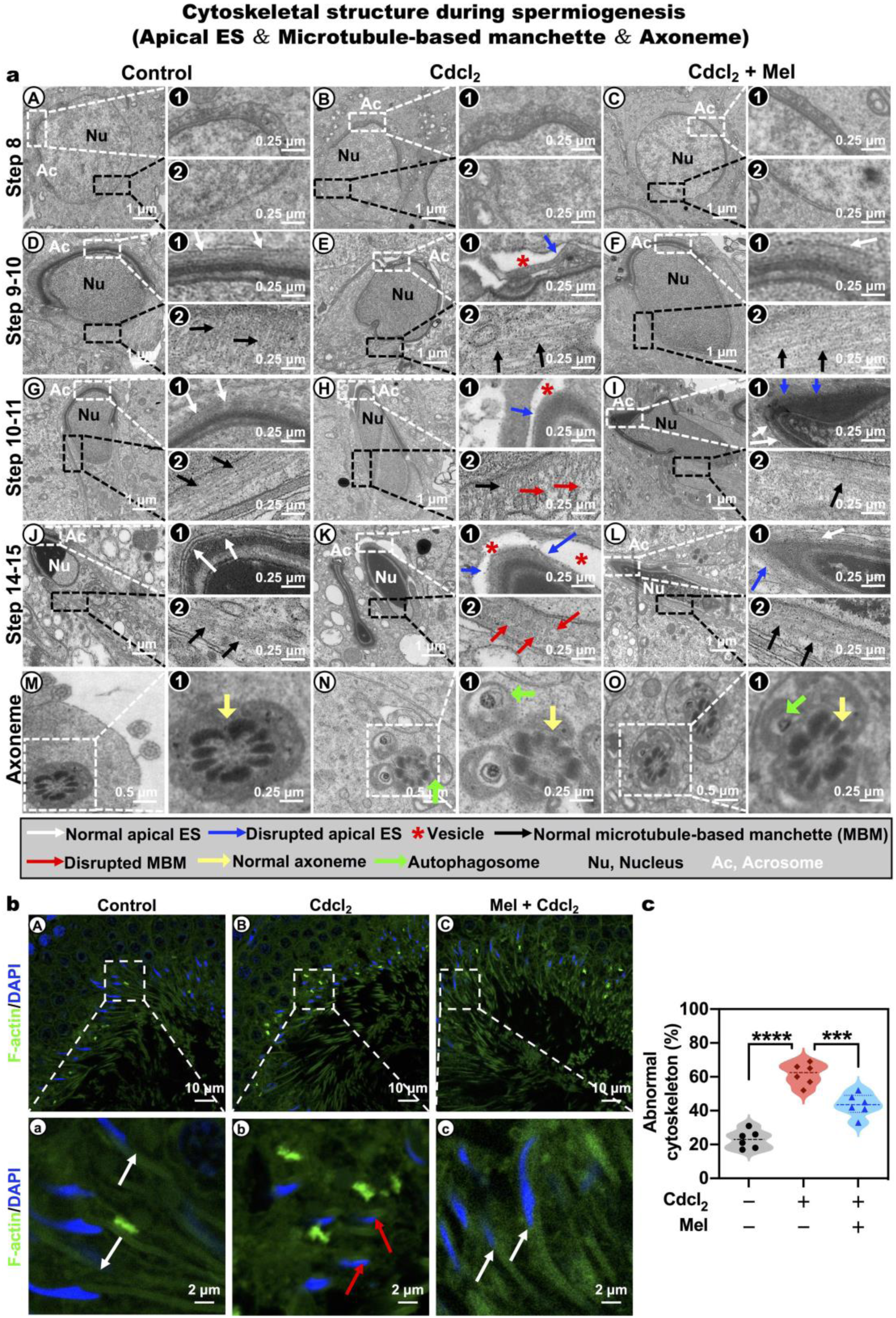
SIRT3 activator Mel rescues spermatid cytoskeleton (apical ES and microtubule-based manchette) assembly disruption induced by Cd during spermiogenesis. **a** Ultrastructural analysis of the spermiogenesis in rat testis during different developmental stages. Areas for the cell (A-O) in the left column (Scale bar, 1 or 0.5 μm) have been shown for further details (1, 2) in the right column (Scale bar, 0.25 μm). (A-C) the ultrastructures of step 8 round spermatids before manchette assembly; (D-F) the ultrastructures of step 9-10 spermatids; (G-I) the ultrastructures of step 10-11 early elongating spermatids spermatids; (J-L) the ultrastructures of step 14-15 late elongating spermatids; (M-O) the ultrastructures of developing axoneme. 1 indicates acrosome, and 2 indicates nuclear membrane in A-C; 1 indicates apical ES, and 2 indicates microtubule-based manchette in D-L. 1 indicates zoom-in axoneme in M-O. Nu, nucleus; Ac, acrosome; white arrowheads indicate normal apical ES; blueberry arrowheads indicate disrupted apical ES; red asterisk indicate vesicle; black arrowheads indicate normal microtubule-based manchette; red arrowheads indicate disrupted microtubule-based manchette; yellow arrowheads indicate normal axoneme; spring green arrowheads indicate autophagosome. **b** Representative confocal microscope imaging of F-actin (a marker of cytoskeleton) in spermatids by the immunofluorescent analysis of phalloidin. Areas (A-C) in the upper panels (Scale bar, 10 μm) have been shown for further details (a-c) in the low panels (Scale bar, 2 μm). White arrowheads indicate normal F-actin structure; red arrowheads indicate disrupted F-actin structure. **c** The percentages of spermatids with abnormal F-actin-containing cytoskeleton. \**p* < 0.05; \*\**p* < 0.01; \*\*\**p* < 0.001; \*\*\*\**p* < 0.0001 in **c**.

F-actin is a marker of cytoskeleton, especially in apical ES and microtubule of the testis. To further confirm the role of SIRT3 activator Mel in the cytoskeleton, we scrutinized the distribution of F-actin in spermatids by the immunofluorescent analysis of phalloidin, which labeled F-actin. Confocal microscopy imaging showed that the F-actin structure was well oriented in linear arrays parallel to the elongating spermatid nucleus’s long axis in the control group (**Fig. 2b (Aa)**). Similar to the TEM experiments, Mel rescued the F-actin structure disruption induced by Cd (**Fig. 2b (Bb, Cc)**), and evidently lowered the percentages of spermatid with abnormal F-actin-containing cytoskeleton (**Fig. 2c**).

Taken together, these results manifested that SIRT3 activator Mel regulated cytoskeleton assembly, including apical ES and microtubule-based manchette in the elongating spermatid during spermiogenesis. Therefore, SIRT3 might participate in cytoskeleton assembly of elongating spermatids during spermiogenesis, influencing the sperm motility.

### Mel regulates SOD2 deacetylation and PDLIM1 by SIRT3 in Cd-induced testicular cytoskeleton assembly disruption

Above TEM and immunofluorescent analysis displayed that Mel modulated cytoskeleton assembly, including BTB with basal ES, apical ES and manchette in the elongating spermatids. To further elucidate the regulatory mechanisms at the molecular level, we asked how Mel modulated cytoskeleton assembly in testis. The protein expression of cytoskeleton assembly markers was investigated. Cd significantly decreased the protein expression of BTB markers—integral membrane proteins (Occludin, JAM-A, N-cadherin), adaptor protein (β-catenin) and regulatory proteins (FAK, p-FAK-Tyr407) (**Fig. 3a, 3b**). In addition to β-catenin, the reduction of other proteins was successfully reversed by Mel. Actually, p-FAK-Tyr407 is a molecular ‘switch’ in the apical ES–BTB axis to regulate BTB restructuring and apical ES adhesion. These results corresponded to the changes of TEM in BTB and apical ES. PDLIM1, NUDC are the identified F-actin networks negative regulator, whose overexpression results in the disassembly of the F-actin net structure in Sertoli cells (Liu et al., 2016). Besides, PDLIM1 is critical for ES assembly. Our results showed that SIRT3 activator Mel reversed the protein expression augment of PDLIM1 and NUDC induced by Cd in testis (**Fig. 3c, 3d**). Therefore, Mel might regulate cytoskeleton assembly by PDLIM1.

**Fig. 3.**
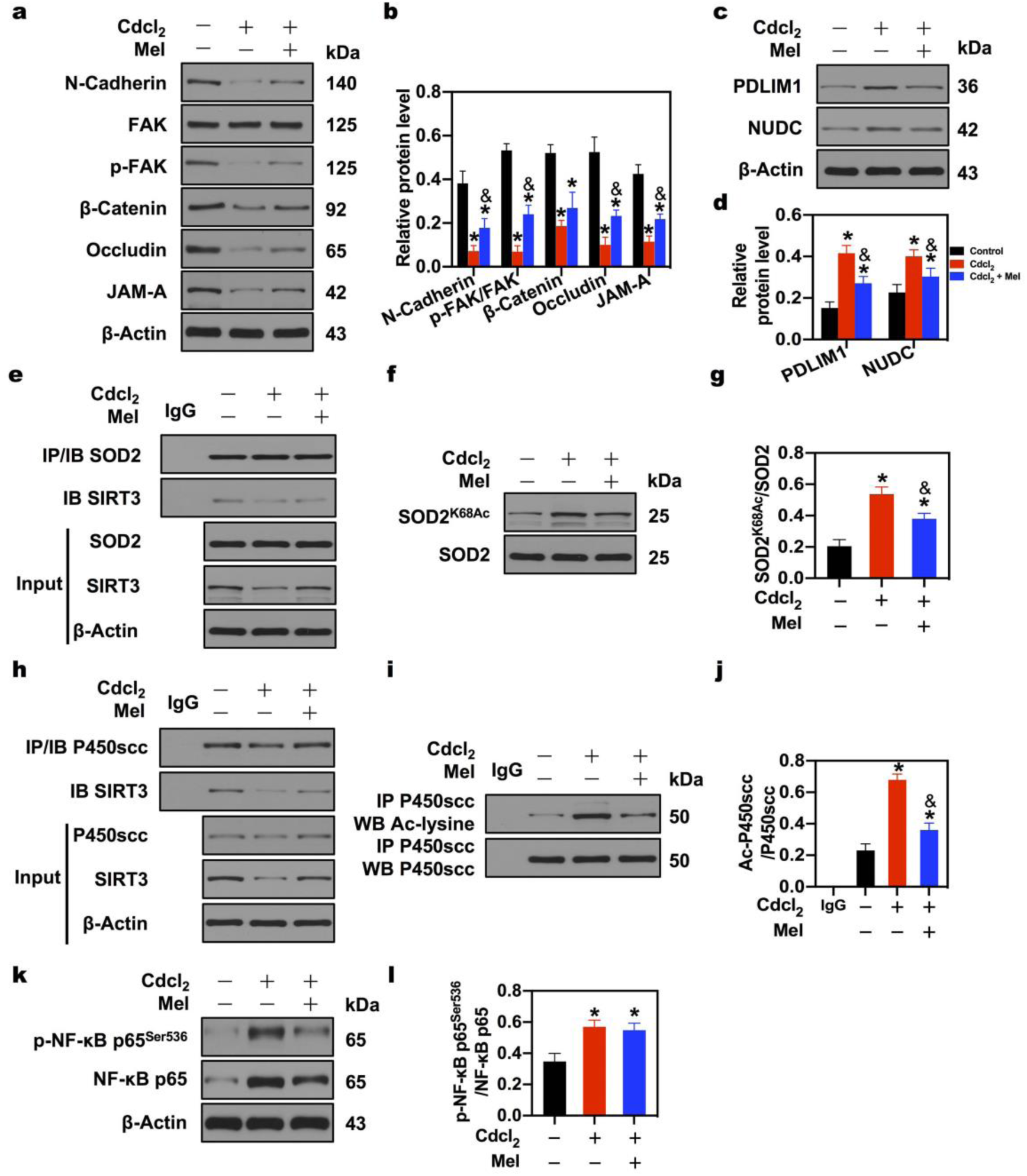
Mel regulates SOD2/P450scc deacetylation by SIRT3, and PDLIM1, NF-κB p65^Ser536^ phosphorylation and P450scc deacetylation by SIRT3 in Cd-induced testicular injury. **a-b** A representative immunoblot and quantification analysis of BTB markers N-Cadherin, β-Catenin, Occludin, JAM-A and apical ES–BTB axis switch FAK, p-FAK-Tyr407 in testis. **c-d** A representative immunoblot and quantification analysis of cytoskeleton negative regulators PDLIM1, NUDC in testis. **e** Co-immunoprecipitation (Co-IP) of SOD2 and SIRT3 in testis. **f-g** A representative immunoblot and quantification analysis of SOD2 acetylation level. **h** Co-IP of P450scc and SIRT3 in testis. **i-j** IP of P450scc and Ac-lysine, and quantification analysis of P450scc acetylation level in testis. **k-l** A representative immunoblot and quantification analysis of NF-κB p65^Ser536^ phosphorylation. \**p* < 0.05 compared with the control. &*p* < 0.05 compared with the Cd-treated group.

Our previous studies showed that Cd elevated testicular ROS level and reduced SOD activity in testis (Chen et al., 2018a) and serum (Wang et al., 2020). SIRT3 is the most robust mitochondrial deacetylase, and SOD2 (also named MnSOD) is just the identified downstream of SIRT3 in the liver (Tao et al., 2010). To investigate the correlation of SIRT3 and SOD2 in testis, we examined the interaction of SIRT3 and SOD2 by co-immunoprecipitation (co-IP). The results indicated that Cd decreased the protein expression of SIRT3 and disturbed the interaction of SIRT3 and SOD2; whereas, SIRT3 activator Mel could effectively increase the protein expression of SIRT3 and the affinity of SIRT3 and SOD2 (**Fig. 3e**). Notably, Cd or Mel had no significant effect on the protein expression of SOD2 (**Fig. 3e**). Then, what is the direct mechanism of the interaction between SIRT3 and SOD2? Whether does SIRT3 exert its role as deacetylase?

To test the hypothesis, SOD2 acetylation levels were determined by the anti-SOD2 antibody with anti-SOD2^K68Ac^ antibody. Significantly, Cd elevated the acetylation of SOD2; Mel reduced the acetylation of SOD2 compared with the Cd-treated group (**Fig. 3f, 3g**), suggesting that SIRT3 regulated SOD2 deacetylation by directly interacting with SOD2 to mediate oxidative stress in testis.

These data indicated that Mel regulated SOD2 deacetylation and PDLIM1 by SIRT3 in Cd-induced testicular cytoskeleton assembly disruption.

### Mel regulates P450scc deacetylation by SIRT3 and NF-κB p65^Ser536^ phosphorylation in Cd-induced testicular cholesterol metabolism disruption

In cholesterol metabolism, cholesterol is delivered into the mitochondria through StAR (gene name Star), and converted to pregnenolone by the catalysis of P450scc (Hsu et al., 2006), which has been considered the rate-limiting step of testosterone biosynthesis. Although Cd increased the expression of Star and cholesterol level, the testosterone level is decreased (**Fig. 1**). A study proposed that the deacetylation of P450scc was associated with SIRT3 and SIRT5 in resveratrol-mediated cortisol biosynthesis (Li et al., 2012). Thus, we speculated whether Mel operated P450scc deacetylation via SIRT3 in this process. Co-IP was performed to verify the interaction of SIRT3 and P450scc. Cd decreased the protein expression of SIRT3 and disturbed the interaction of SIRT3 and P450scc; whereas, Mel effectively increased the protein expression of SIRT3 and the affinity of SIRT3 and P450scc (**Fig. 3h**). Whether Cd or Mel had no significant effect on the protein expression of P450scc (**Fig. 3h**). Synchronously, P450scc acetylation levels were determined by immunoprecipitation with an anti-P450scc antibody, followed by immunoblot analysis of acetylated-lysine. Cd elevated the acetylation of P450scc; Mel reduced the acetylation of P450scc compared with the Cd-treated group (**Fig. 3i, 3j**), suggesting that SIRT3 regulated P450scc deacetylation by directly interacting with P450scc to mediate cholesterol metabolism in testis.

Then, how does Mel exert its role on SIRT3 in testis? Given that Mel could remarkably alleviate inflammasome-induced pyroptosis by blocking NF-κB p65 signal in mice adipose tissue (Liu et al., 2017b), and the inhibition of NF-κB p65 regulated testosterone production by Nur77 and SF-1 (Hong et al., 2004), thus the activity of NF-κB p65 was observed. We noticed that Cd indeed stimulated NF-κB p65^Ser536^ phosphorylation, and Mel mitigated the phosphorylation of NF-κB p65^Ser536^. Meanwhile, Mel reversed Cd-induced protein expression changes of NF-κB p65 and p-NF-κB p65^Ser536^ respectively. Nonetheless, the ratio of p-NF-κB p65^Ser536^/NF-κB p65 did not exhibit a significant difference between the Cd-treated group and Cd with Mel group (**Fig. 3k, 3l**). Above results evidenced that Mel regulates P450scc deacetylation by SIRT3 and NF-κB p65 in Cd-induced testicular cholesterol metabolism disruption.

### Mel-mediated cholesterol metabolism protection is dependent on Sirt3, which is the upstream regulator of P450scc deacetylation, but not of NF-κB p65^ser536^ phosphorylation in TM3 mouse Leydig cells

To further validate the role of SIRT3 in cholesterol metabolism in Leydig cells, we generated Sirt3 overexpression and knockdown models in TM3 mouse Leydig cells (TM3 cells). The efficiencies of Sirt3 overexpression and knockdown were examined (**supplementary Fig. 1c**).

Initially, we got the half-maximal inhibitory concentration (IC50) of Cd for TM3 cells—8.725 μg/ml by the concentration gradient method (Wang et al., 2020), which was exploited in subsequent experiments. Cell Counting Kit-8 (CCK-8) assay indicated that Mel significantly rescued cell viability reduction induced by Cd (**Fig. 4a_1_**—the first panel in Fig. 4a), implying that Mel blocked the specific injury process. To determine whether Mel-mediated protection depends on SIRT3 or not, Sirt3 overexpression adenovirus (Ad-Sirt3) and Sirt3 knockdown adenovirus with shRNA (Sh-Sirt3) were utilized. Like Mel, the overexpression of Sirt3 rescued cell viability reduction induced by Cd (**Fig. 4a_2_**), suggesting that Sirt3 participated in the specific protection process. Interestingly, the knockdown of Sirt3 decreased TM3 cell viability, but Mel failed to rescue sh-Sirt3-induced cell viability reduction (**Fig. 4a_3_**), revealing that Sirt3 dominated the protection process, and scilicet Mel-mediated cell viability protection was dependent on Sirt3 in TM3 cells.

**Fig. 4.**
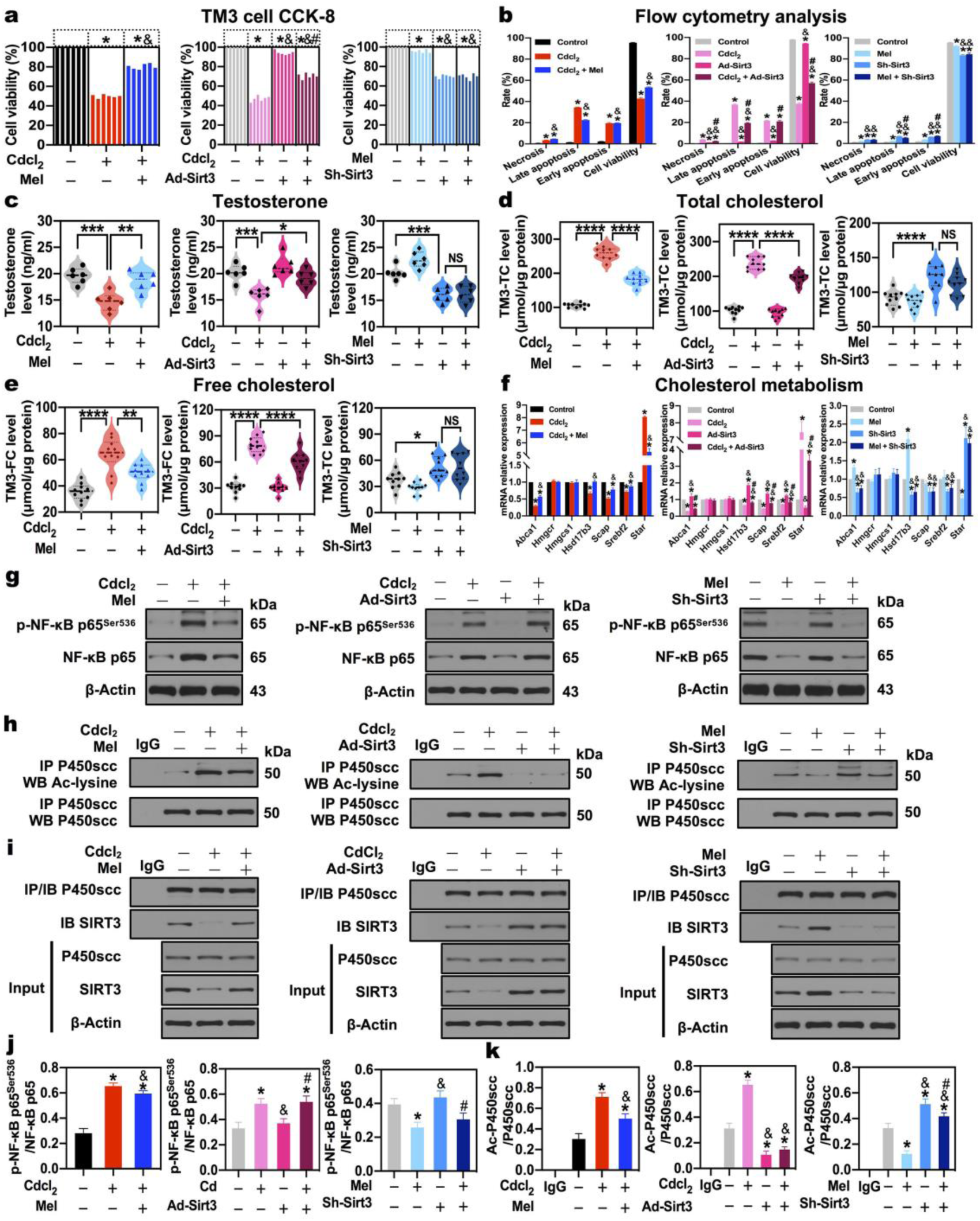
Mel-mediated cholesterol metabolism protection is dependent on Sirt3, which is the upstream regulator of P450scc deacetylation, but not of NF-κB p65^ser536^ phosphorylation in TM3 mouse Leydig cells. **a** Cell viability analysis by CCK-8 assay. **b** Necrosis, late apoptosis, early apoptosis, cell viability analysis by flow cytometry. **c** Testosterone level in TM3 cells supernatants. **d** Total cholesterol level in TM3 cells. **e** Free cholesterol level in TM3 cells. **f** mRNA expression levels of cholesterol metabolism markers of Abca1, Hmgcr, Hmgcs1, Hsd17b1, Scap, Srebf2, Star in TM3 cells. **g, j** A representative immunoblot and quantification analysis of NF-κB p65^Ser536^ phosphorylation in TM3 cells. **h, k** IP of P450scc and Ac-lysine, and quantification analysis of P450scc acetylation level in TM3 cells. **i** Co-IP of P450scc and SIRT3 in TM3 cells. NS, *p* > 0.05; \**p* < 0.05; \*\**p* < 0.01; \*\*\**p* < 0.001; \*\*\*\**p* < 0.0001 in **a-e**. \**p* < 0.05 compared with the first group (control). &*p* < 0.05 compared with the second group, #*p* < 0.05 compared with the third group in each panel of **f, j, k**.

Flow cytometry analysis confirmed the results of the CCK-8 assay. Cd significantly decreased the percentage of viable cells and increased the percentage of necrotic, early and late apoptotic cells; remarkably, Mel reversed Cd-triggered TM3 cell apoptosis (**Supplementary Fig. 2a; Fig. 4b_1_**). Moreover, the overexpression of Sirt3 rescued apoptosis induced by Cd, but Mel did not rescue sh-Sirt3-induced apoptosis in TM3 cells (**Supplementary Fig. 2b, 2c; Fig. 4b_2,3_**), demonstrating that Mel-mediated cell apoptosis protection was dependent on Sirt3 in TM3 cells.

**Supplementary Fig. 1.**
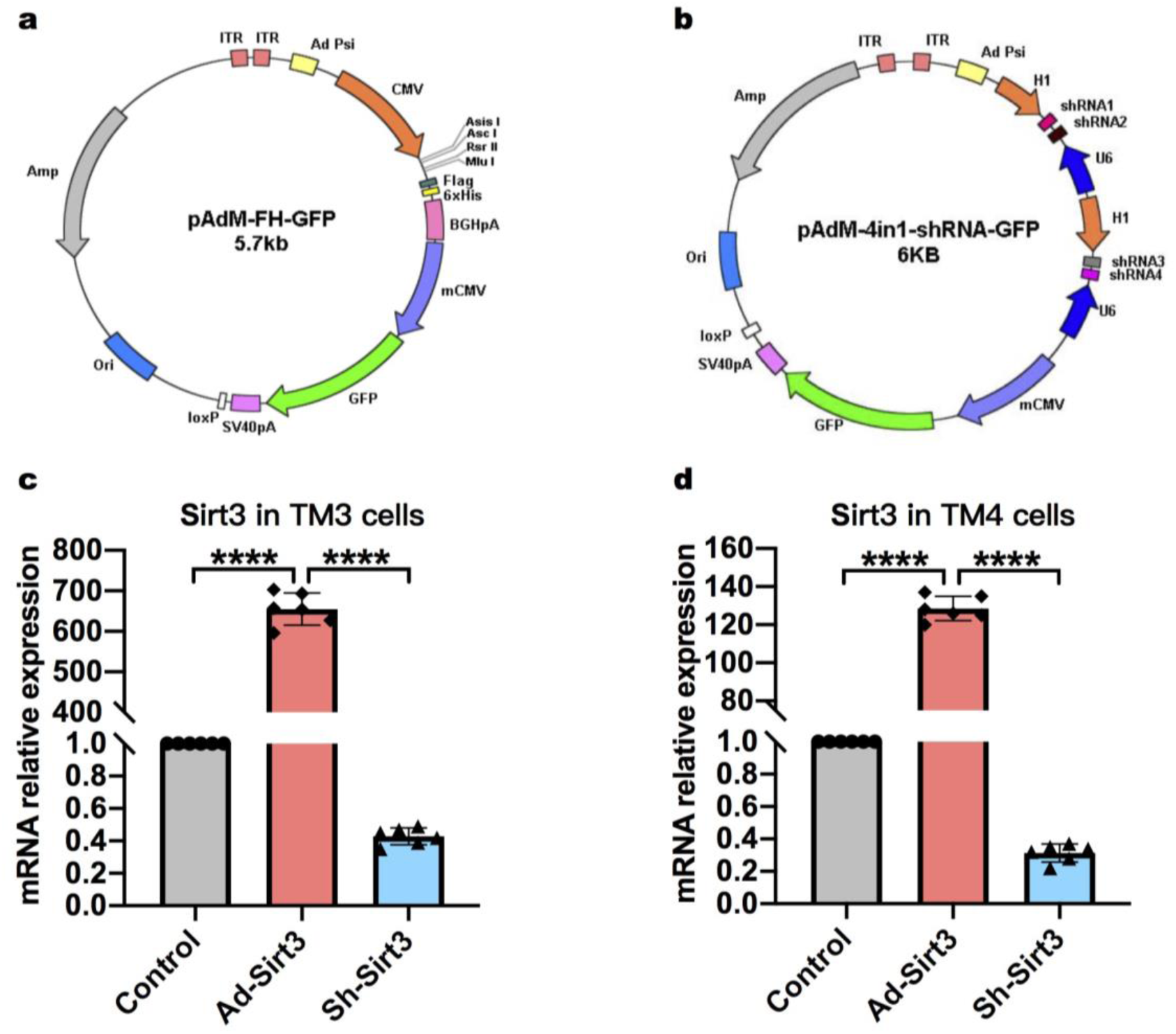
The efficiency of Sirt3 overexpression or knockdown in TM3 cells and TM4 cells. **a** The adenovirus vector for Sirt3 overexpression. **b** The shRNA-containing adenovirus vector for Sirt3 knockdown. **c** The efficiency of Sirt3 overexpression and knockdown in TM3 cells by mRNA expression level analysis. **d** The efficiency of Sirt3 overexpression and knockdown in TM4 cells by mRNA expression level analysis. \*\*\*\**p* < 0.0001.

**Supplementary Fig. 2.**
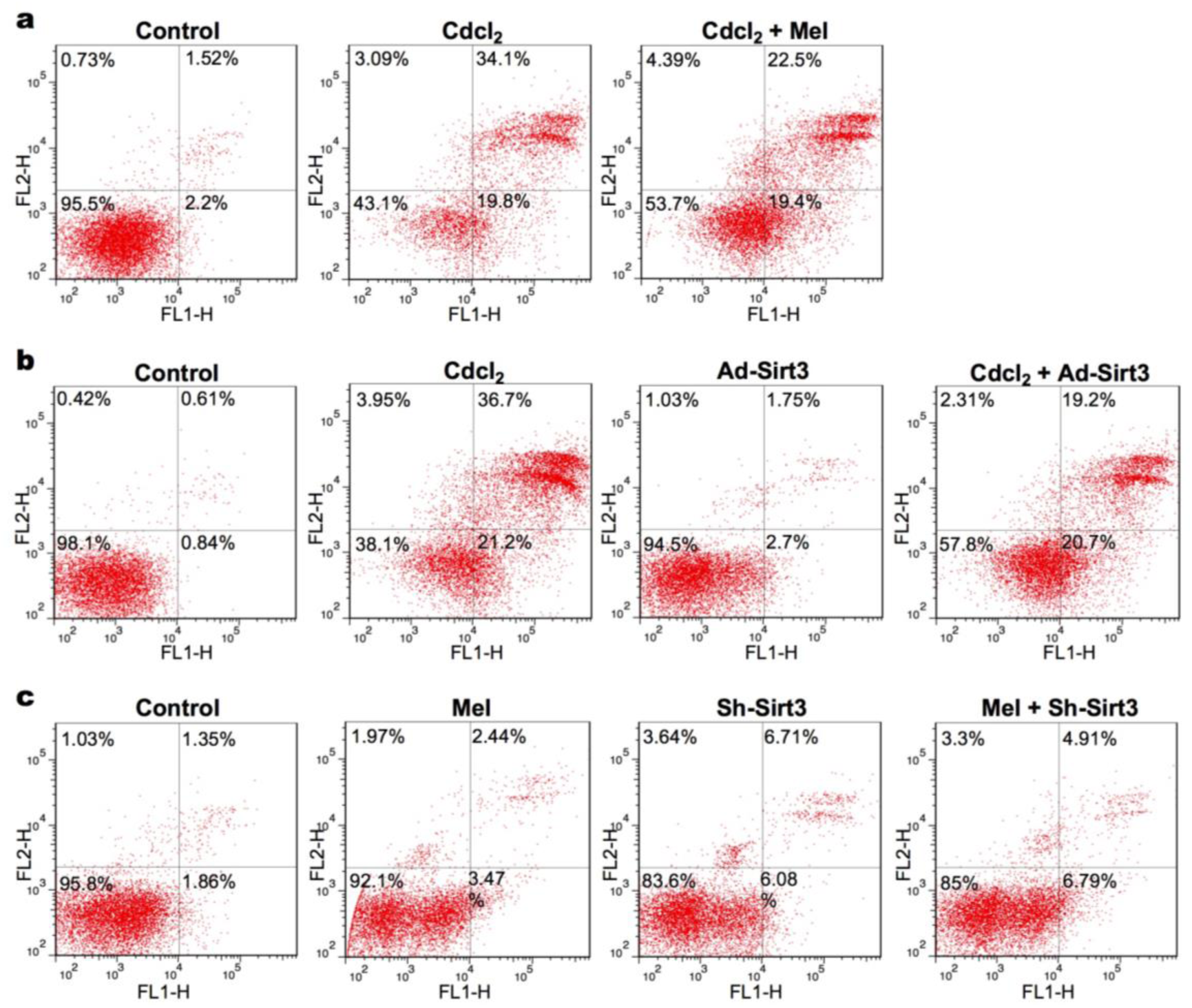
Mel-mediated cell apoptosis protection is dependent on Sirt3 in TM3 cells. Representative flow cytometry analysis: **a** Mel reverses Cd-triggered TM3 cell apoptosis. **b** The overexpression of Sirt3 rescues apoptosis induced by Cd in TM3 cells. **c** Mel fails to rescue sh-Sirt3-induced apoptosis in TM3 cells.

Subsequently, the level of testosterone, total cholesterol (TC), and free cholesterol (FC) in TM3 cells were scrutinized. Consistent with in vivo results, Cd decreased testosterone level and increased TC and FC level, which were reversed by Mel (**Fig. 4c_1_, 4d_1_, 4e_1_**). Likewise, the overexpression of Sirt3 defended testosterone, TC, and FC disruption induced by Cd in TM3 cells (**Fig. 4c_2_, 4d_2_, 4e_2_**). The knockdown of Sirt3 led to analogous testosterone, TC, and FC disruption as Cd, which did not be salvaged by Mel (**Fig. 4c_3_, 4d_3_, 4e_3_**), indicating that Mel-mediated testosterone and cholesterol protection was dependent on Sirt3 in TM3 cells.

Testosterone synthesis is derived from the cholesterol mechanism. To further clarify, in Leydig cell, the specific effect of SIRT3 on regulating testosterone and cholesterol, we detected cholesterol metabolism markers, including Hmgcr and Hmgcs1 (cholesterol biosynthesis markers), Abca1 (cholesterol efflux marker), Star (mitochondrial cholesterol transporter), Scap/Srebf2 (cholesterol metabolism regulators), Hsd17b3 (testosterone biosynthesis marker). Both Hmgcr and Hmgcs1 were not disturbed by Cd, Mel, Ad-Sirt3 and sh-Sirt3, suggesting that Sirt3 did not correlate with cholesterol biosynthesis (**Fig. 4f**). Cd significantly diminished the expression of Abca1 and elevated the expression of Star, implying that Cd enhanced cholesterol influx; Cd reduced the expression of Scap, Srebf2 and Hsd17b3, implying that Cd impaired the regulation of cholesterol metabolism; however, Mel rescued cholesterol metabolism disruption by reversing the expression of Abca1, Star, Scap, Srebf2 and Hsd17b3 (**Fig. 4f_1_**). In parallel, the overexpression of Sirt3 protected from cholesterol metabolism disruption induced by Cd in TM3 cells (**Fig. 4f_2_**). Instead, the knockdown of Sirt3 induced cholesterol metabolism disruption similar to Cd, which did not be rescued by Mel (**Fig. 4f_3_**), indicating that Mel-mediated cholesterol metabolism was dependent on Sirt3 in TM3 cells.

In vivo study, we found Mel might regulate NF-κB p65^ser536^ phosphorylation to mediate SIRT3 with P450scc deacetylation in Cd-induced testicular cholesterol metabolism disruption. Herein, NF-κB p65^ser536^ phosphorylation and P450scc deacetylation were investigated in TM3 mouse Leydig cells. As expected, Cd significantly stimulated NF-κB p65^Ser536^ phosphorylation, which was weakened by Mel in TM3 cells (**Fig. 4g_1_, 4j_1_**). Nonetheless, the overexpression of Sirt3 showed no difference in NF-κB p65^Ser536^ phosphorylation compared with the control group and failed to reverse Cd-stimulated NF-κB p65^Ser536^ phosphorylation (**Fig. 4g_2_, 4j_2_**). Despite that Mel indeed inhibited NF-κB p65^Ser536^ phosphorylation, the knockdown of Sirt3 didn’t influence NF-κB p65^Ser536^ phosphorylation (**Fig. 4g_3_, 4j_3_**). Results suggested that, in TM3 cells, Mel regulated NF-κB p65^Ser536^ phosphorylation, which was not dominated by Sirt3.

Subsequently, we found that Cd promoted P450scc acetylation, which was rescued by Mel in TM3 cells (**Fig. 4h_1_, 4k_1_**). Noticeably, the overexpression of Sirt3 lessened P450scc acetylation and reversed Cd-induced P450scc acetylation (**Fig. 4h_2_, 4k_2_**); the knockdown of Sirt3 dramatically boosted P450scc acetylation and reversed Mel-induced P450scc deacetylation (**Fig. 4h_3_, 4k_3_**), indicating that Mel regulated P450scc deacetylation, which was dominated by Sirt3.

To further confirm the interaction between SIRT3 and P450scc, co-IP experiment was performed in TM3 cells. Consequently, Cd, Mel, Ad-Sirt3 or sh-Sirt3 had no significant effect on the protein expression of P450scc; strikingly, Cd reduced the interaction of SIRT3 and P450scc, which was reversed by Mel (**Fig. 4i_1_**). The overexpression of Sirt3 contributed to the interaction of SIRT3 and P450scc and rescued Cd-induced disruption (**Fig. 4i_2_**); the knockdown of Sirt3 impaired the interaction of SIRT3 and P450scc, which failed to be reversed by Mel (**Fig. 4i_3_**), displaying that SIRT3 directly interacted with P450scc to mediate P450scc deacetylation.

Above results demonstrated that Mel-mediated cholesterol metabolism protection was dependent on Sirt3, which was the upstream regulator of P450scc deacetylation, but not of NF-κB p65^ser536^ phosphorylation in TM3 mouse Leydig cells.

### NF-κB, as the upstream transcription factor, represses Sirt3 transcription by binding to TFBS1 and TFBS2 of Sirt3 promoter in TM3 mouse Leydig cells

Due to three facts as follows: (1) Mel-mediated cholesterol metabolism was dependent on Sirt3 in TM3 cells; (2) Mel regulated NF-κB p65^Ser536^ phosphorylation in TM3 cells; (3) Sirt3 was not the upstream regulator of NF-κB p65^ser536^ phosphorylation in TM3 cells, we speculated that Sirt3 was the downstream of NF-κB p65 in Mel-mediated cholesterol metabolism. Phosphorylation of p65 at Ser536 led to nuclear localization of the transcriptionally active complex, and NF-κB mediated transactivation of several downstream genes (Mai et al., 2020). To test the conjecture, we examined the cytoplasmic and nuclear localization of NF-κB molecules in Cd with or without Mel-treated TM3 cells. As shown in **Fig. 5a**, Cd resulted in slightly increased nuclear translocation of c-Rel and robust nuclear translocation of p50, p65 and RelB, which could be ameliorated by Mel, suggesting that phosphorylation of p65 at Ser536 indeed stimulated nuclear translocation of NF-κB molecules in TM3 cells.

**Fig. 5.**
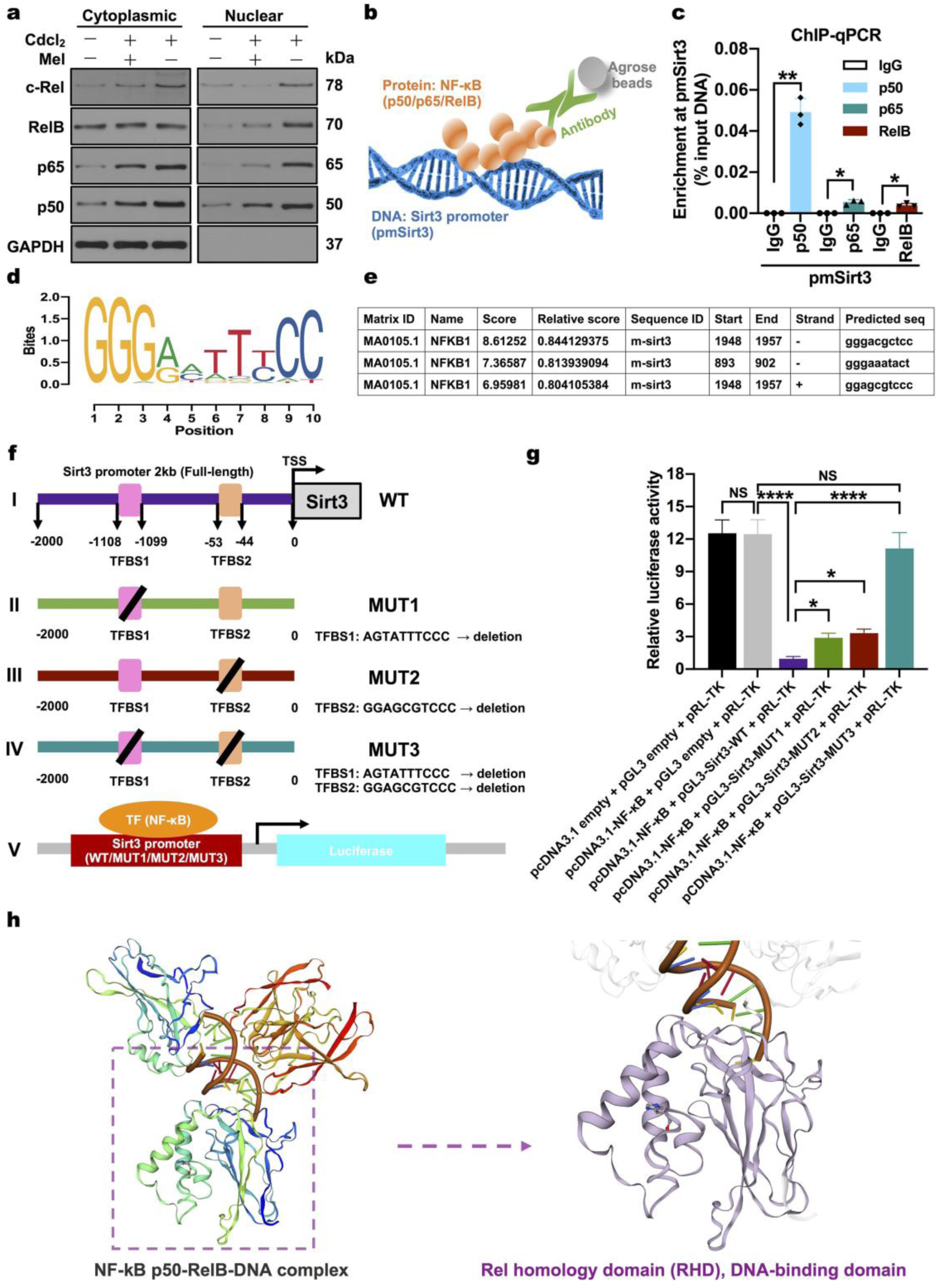
NF-κB, as upstream transcription factor, represses Sirt3 transcription by binding to TFBS1 and TFBS2 of Sirt3 promoter in TM3 mouse Leydig cells. **a** Western blot analysis of the cytoplasmic and nuclear localization of NF-κB molecules p50, p65, RelB, c-Rel. **b** ChIP assay model for NF-κB molecules p50, p65, RelB and Sirt3 promoter. **c** ChIP assay of p50, p65 and RelB, especially p50, binding to Sirt3 promoter in TM3 cells. **d** Position weight matrix (PWM) and motif logo of NF-κB1 p50 TF–DNA binding preferences. **e** Predicted transcription factor binding sites (TFBSs) with a relative profile score of greater than 0.8 by JASPAR database. **f** A diagram showing the relative positions of full-length (FL) and predicted TFBSs of Sirt3 promoter reporters and wild-type (WT) and mutant promoter constructions followed by dual-luciferase reporter assay. I the relative positions of FL and predicted TFBSs of Sirt3 promoter (WT); II deletion of TFBS1 (−1108 to −1099, AGTATTTCCC) for constructing mutant Sirt3 promoter 1 (MUT1); III deletion of TFBS2 (−53 to −44, GGAGCGTCCC) for constructing mutant Sirt3 promoter 2 (MUT2); IV simultaneous deletion of TFBS1 and TFBS2 for constructing mutant Sirt3 promoter 3 (MUT3); V a schematic of dual-luciferase reporter assay for detecting the interaction between NF-κB1 p50 and Sirt3 promoter (WT/MUT1/MUT2/MUT3). **g** Dual-luciferase reporter assay results showing NF-κB1 p50-dependent suppression of Sirt3 transcription by simultaneously binding to TFBS1 and TFBS2 in Sirt3 promoter in HEK293T cells. **h** 3D structure of NF-kB p50-RelB-DNA complex from Swiss-Model. Left panel, NF-kB p50-RelB-DNA complex; right panel, Rel homology domain (RHD), DNA-binding domain. NS, *p* > 0.05; \**p* < 0.05; \*\**p* < 0.01; \*\*\**p* < 0.001; \*\*\*\**p* < 0.0001.

Querying UCSC Genome browser (https://genome.ucsc.edu) with expanded JASPAR database, NF-κB subunits, including RELA (also named p65), RELB and REL, were predicted as putative upstream transcription factors for mouse Sirt3 gene (**Supplementary Fig. 3**). To confirm the interaction of NF-κB and Sirt3 promoter region in TM3 mouse Leydig cells, chromatin immunoprecipitation (ChIP) assay was performed as described in **Fig. 5b**. Protein-DNA complexes were extracted from TM3 cells and immunoprecipitated using anti-NF-κB (p50, p65, RelB respectively) with precipitation of normal IgG as the negative control. DNA fragments were amplified with primers specific for the Sirt3 promoter sequence. ChIP-qPCR assay manifested a physiological binding of p65, RelB and especially p50 to the Sirt3 promoter in TM3 cells (**Fig. 5c**).

**Supplementary Fig. 3.**
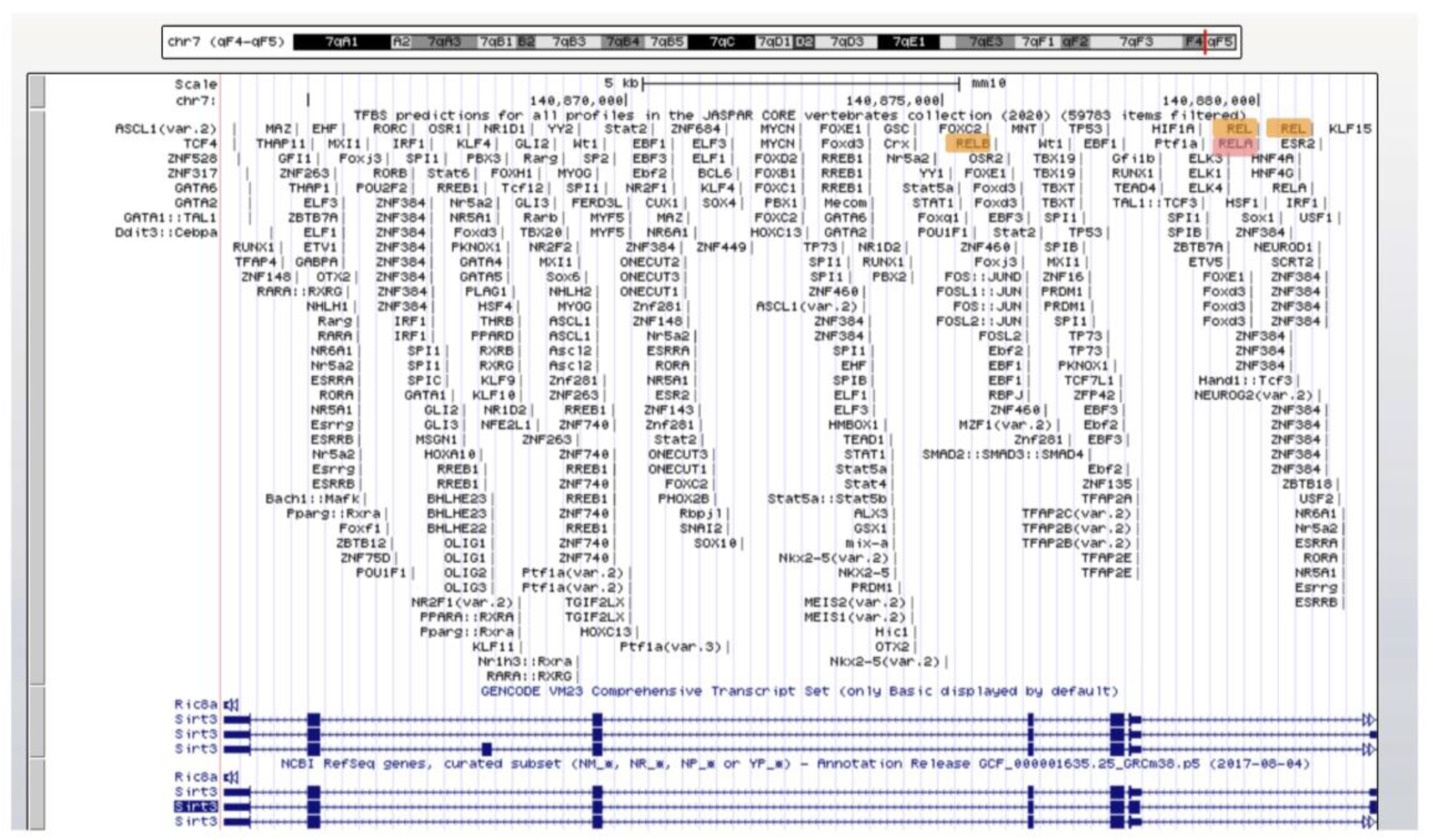
Prediction of upstream transcription factors for mouse Sirt3 gene. By Querying UCSC Genome browser (https://genome.ucsc.edu) with expanded JASPAR database, NF-κB subunits, including RELA (also named p65), RELB and REL, were predicted as putative upstream transcription factors for mouse Sirt3 gene.

To further substantiate the transcriptional regulation of Sirt3 by NF-κB and recognize the specific transcription factor (TF)-DNA binding sequences, we identified the consensus sequence of mouse NF-κB p50 by analyzing structural profiles of transcription factor binding sites (TFBSs) from JASPAR and UniPROBE databases. In fact, the consensus sequence reflected p50 TF–DNA binding preferences, which were represented by a position weight matrix (PWM) and visualized as motif logo (**Fig. 5d**). Based on the selected matrix models, three fragments in full-length mouse Sirt3 promoter (2kb) were predicted as TFBSs with a relative profile score of greater than 0.8 by the JASPAR database (**Fig. 5e**). Given that gggacgctcc on reverse (minus) DNA strand (1948-1957) overlapped with ggagcgtccc on forward (plus) DNA strand (1948-1957), there were actually two fragments as putative TFBS1 and TFBS2. In the full-length (FL) Sirt3 promoter from −2000 bp to 0 transcription start site (TSS), TFBS1 (AGTATTTCCC) locates in −1108 to −1099, and TFBS2 (GGAGCGTCCC) locates in −53 to −44 (**Fig. 5f(I)**). Wild-type (WT) or TFBS1/TFBS2 mutant Sirt3 promoter (MUT1/MUT2/MUT3) luciferase reporter vector was transiently co-transfected with NF-κB p50 overexpression plasmid (pcDNA3.1-NF-κB) into HEK 293T cells (**Fig. 5f**), and relative luciferase activity was measured as a function of NF-κB p50-dependent Sirt3 transcription. Conspicuously, NF-κB p50 with WT-Sirt3 promoter inhibited the transcription, while NF-κB p50 with TFBS1/TFBS2 mutant Sirt3 promoter (MUT1/MUT2/MUT3) attenuated the transcriptional suppression; especially, MUT3 (simultaneous deletion of TFBS1 and TFBS2) had no difference in transcription compared with empty pGL3-promoter vector (**Fig. 5g**). Results demonstrated that NF-κB p50 could suppress Sirt3 transcription by binding to TFBS1 and TFBS2 in Sirt3 promoter (**Fig. 5h**), which responded to the decreased protein expression of SIRT3 in Cd-treated testis and TM3 cells.

Above stringing shreds of evidence supported the inference that, **in Leydig cells, phosphorylation of NF-κB p65^Ser536^ stimulated the nuclear translocation of NF-κB molecules (subunits p50, p65, RelB), which bound to the promoter of Sirt3; NF-κB (particularly p50) repressed Sirt3 transcription by binding to TFBS1 and TFBS2 in Sirt3 promoter; then, the deficiency of SIRT3 disturbed cholesterol metabolism and testosterone synthesis by impairing P450scc deacetylation in male subfertility model induced by Cd.**

### Overexpression and knockdown of SIRT3 blunts and enhances PDLIM1 and SOD2 acetylation in Mel-mediated TM4 mouse Sertoli cell protection

Except for the role of Sirt3 in cholesterol metabolism of Leydig cell, our in vivo study displayed that Mel might regulate SOD2 deacetylation and PDLIM1 by SIRT3 in Cd-induced the disruption of cytoskeleton assembly, including basal ES (Sertoli-Sertoli cell interface), apical ES (Sertoli-elongating spermatid interface) and microtubule-based manchette. Considering that the disturbed cytoskeleton was related to the Sertoli cell, and PDLIM1 in Sertoli cell could regulate ES (Liu et al., 2016), we hypothesized that SIRT3 orchestrated cytoskeleton assembly by PDLIM1 and SOD2 deacetylation in Sertoli cell.

To verify the hypothesis, Sirt3 overexpression and knockdown models in TM4 mouse Sertoli cells (TM4 cells) were established. The efficiency of Sirt3 overexpression and knockdown were examined (**supplementary Fig. 1d**). IC50 of Cd for TM4 cells—12 μg/ml by the concentration gradient method (Wang et al., 2020) was exploited in subsequent experiments. Similar to TM3 cells, CCK-8 assay indicated that both Mel and overexpression of Sirt3 rescued TM4 cell viability reduction induced by Cd (**Fig. 6a_1,2_**); the knockdown of Sirt3 decreased cell viability, but Mel failed to rescue sh-Sirt3-induced cell viability reduction (**Fig. 6a_3_**), hinting that Sirt3 dominated the protection process. Scilicet Mel-mediated cell viability protection was dependent on Sirt3 in TM4 cells.

**Fig. 6.**
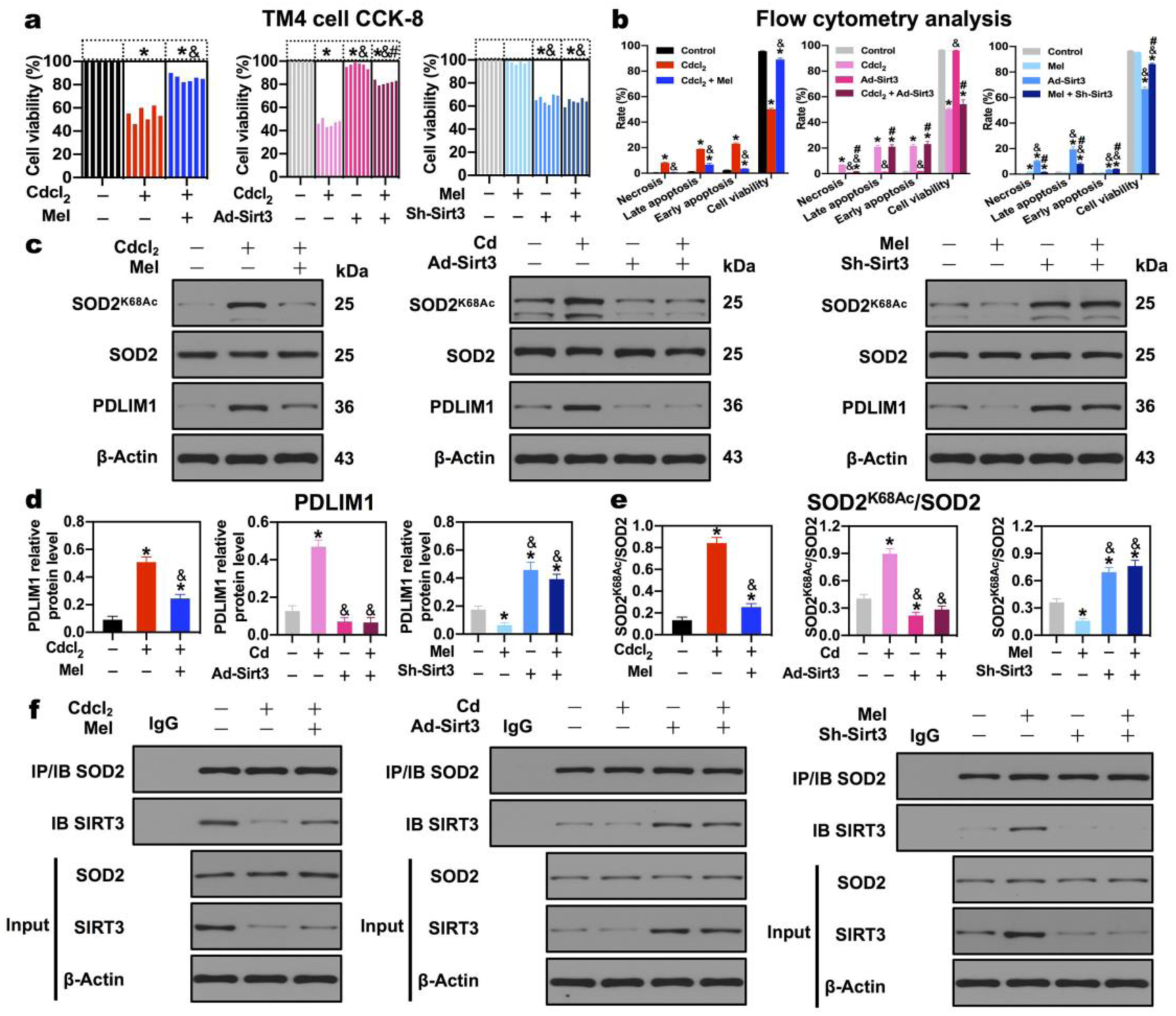
Overexpression and knockdown of SIRT3 blunts and enhances PDLIM1 and SOD2 acetylation in Mel-mediated TM4 mouse Sertoli cell protection. **a** Cell viability analysis by CCK-8 assay. **b** Necrosis, late apoptosis, early apoptosis, cell viability analysis by flow cytometry. **c** A representative immunoblot of cytoskeleton negative regulator PDLIM1, SOD2 and SOD2^K68Ac^ in TM4 cells. **d** Quantification analysis of cytoskeleton negative regulator PDLIM1 protein level in TM4 cells. **e** Quantification analysis of SOD2 acetylation level in TM4 cells. **f** Co-IP of SOD2 and SIRT3 in TM4 cells. \**p* < 0.05 compared with the first group (control). &*p* < 0.05 compared with the second group, #*p* < 0.05 compared with the third group in each panel of **a, b, d, e**.

Flow cytometry analysis detected the cell viability and apoptosis. Cd significantly decreased the percentage of viable cells and increased the percentage of necrotic, early and late apoptotic cells; whereas, Mel reversed Cd-triggered TM4 cell apoptosis (**Supplementary Fig. 4a; Fig. 6b_1_**). The overexpression of Sirt3 rescued necrosis but not apoptosis in Cd-treated TM4 cells; the knockdown of Sirt3 facilitated apoptosis, which couldn’t be attenuated by Mel in TM4 cells (**Supplementary Fig. 4b, 4c; Fig. 6b_2,3_**), proving that Mel-mediated cell apoptosis protection was dependent on Sirt3 in TM4 cells.

**Supplementary Fig. 4.**
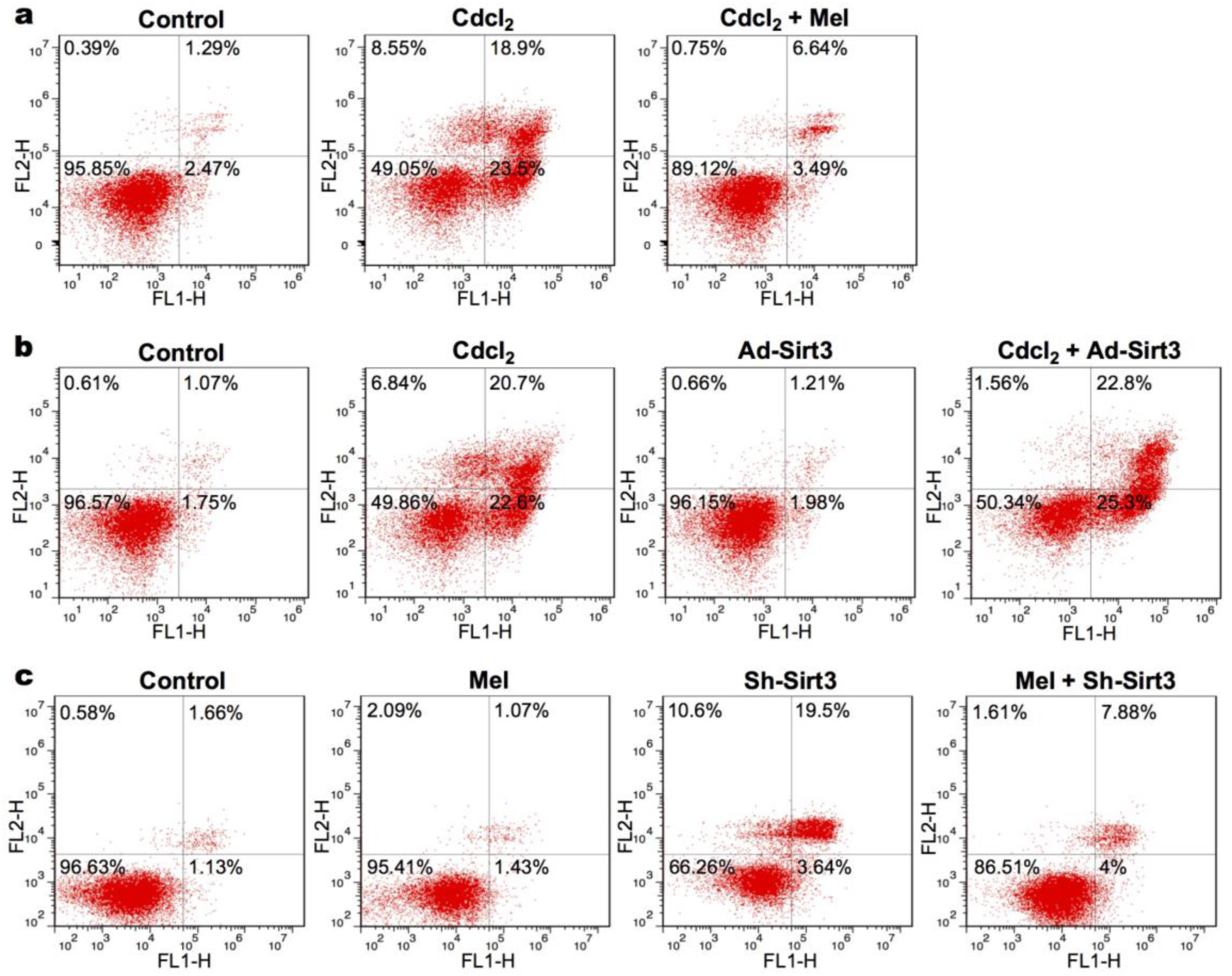
Mel-mediated cell apoptosis protection is dependent on Sirt3 in TM4 cells. Representative flow cytometry analysis: **a** Mel reverses Cd-triggered TM4 cell apoptosis. **b** The overexpression of Sirt3 rescues necrosis but not apoptosis in Cd-treated TM4 cells. **c** Mel fails to rescue sh-Sirt3-induced apoptosis in TM4 cells.

To address whether SIRT3 regulated SOD2 deacetylation and PDLIM1 in Sertoli cells, we assessed the PDLIM1 protein levels and SOD2 acetylation when Sirt3 was overexpressed or restrained. PDLIM1, as negative cytoskeleton regulator, was ignited in Cd-treated TM4 cells; whereas, Mel extinguished the augment of PDLIM1 protein level induced by Cd (**Fig. 6c_1_, 6d_1_**). Likewise, the overexpression of Sirt3 reversed the augment of PDLIM1 protein level induced by Cd (**Fig. 6c_2_, 6d_2_**). Prominently, the knockdown of Sirt3 provoked the protein expression of PDLIM1, which couldn’t be salvaged by Mel (**Fig. 6c_3_, 6d_3_**), intimating that Sirt3 was indispensable for the regulation of PDLIM1 in Mel-mediated TM4 cells protection.

In terms of SOD2 acetylation, both Mel and the overexpression of Sirt3 parallelly reversed Cd-induced SOD2 acetylation (**Fig. 6c_1,2_, 6e_1,2_**). Whereas, the knockdown of Sirt3 impede SOD2 deacetylation and reversed Mel-mediated SOD2 deacetylation (**Fig. 6c_3_, 6e_3_**), suggesting that Sirt3 was imperative for SOD2 deacetylation in Mel-mediated TM4 cells protection.

To further confirm the interaction between SIRT3 and SOD2, co-IP experiment was performed in TM4 cells. Consequently, Cd barricaded the interaction of SIRT3 and SOD2, which was reversed by Mel and the overexpression of Sirt3 (**Fig. 6f_1_, 6f_2_**). The overexpression of Sirt3 significantly facilitated the interaction of SIRT3 and SOD2 (**Fig. 6f_2_**); whereas, the knockdown of Sirt3 impaired the interaction of SIRT3 and SOD2, which failed to be reversed by Mel (**Fig. 6f_3_**). Taken together, these data approved that SIRT3 mediated SOD2 deacetylation by directly interacting with SOD2 in TM4 cells.

### Overexpression and knockdown of Sirt3 protects and impairs the F-actin-containing cytoskeleton assembly in Mel-mediated TM4 mouse Sertoli cells protection

F-actin, as a pivotal component of the cytoskeleton, was proposed to be associated with PDLIM1 (also named CLP36) twenty years ago (Bauer et al., 2000). A recent study has demonstrated that PDLIM1 competitively binds to ACTN4, which disaffiliates from F-actin, hampering F-actin organization (Huang et al., 2020). Critically, PDLIM1 was responsible for ES assembly in Sertoli cell (Liu et al., 2016). It’s rational to deem that SIRT3 regulated PDLIM1 and SOD2 deacetylation to operate cytoskeleton assembly. To authenticate the role of SIRT3 in cytoskeleton assembly, we monitored the F-actin distribution by using rhodamine-phalloidin (red) in Sirt3 overexpression and knockdown models of TM4 mouse Sertoli cells.

As depicted in **Fig. 7a**, in normal TM4 cells, a highly ordered F-actin network wove cytoskeleton to formulate the intact cell; Cd collapsed the F-actin organization, yet Mel ameliorated the F-actin disruption induced by Cd. The percentages of abnormal cytoskeletal Sertoli cells per 200 cells were calculated in 6 independent experiments for each group. Cd increased the rate of abnormal Sertoli cells with severely F-actin disruption, which was rescued by Mel (**Fig. 7b**), implying that Mel protected the cytoskeleton assembly in Sertoli cells. The group with Sirt3 overexpression presented a well-organized F-actin structure and salvaged the F-actin disruption induced by Cd (**Fig. 7c**). Meanwhile, the overexpression of Sirt3 decreased the rate of abnormal F-actin Sertoli cells compared with Cd (**Fig. 7d**), suggesting that Sirt3 participated in the Mel-mediated cytoskeleton assembly. Furthermore, the knockdown of Sirt3 apparently disturbed the F-actin network, which couldn’t be rescued by Mel (**Fig. 7e, 7f**), revealing that Mel-mediated cytoskeleton assembly protection was dependent on Sirt3 in TM4 cells.

**Fig. 7.**
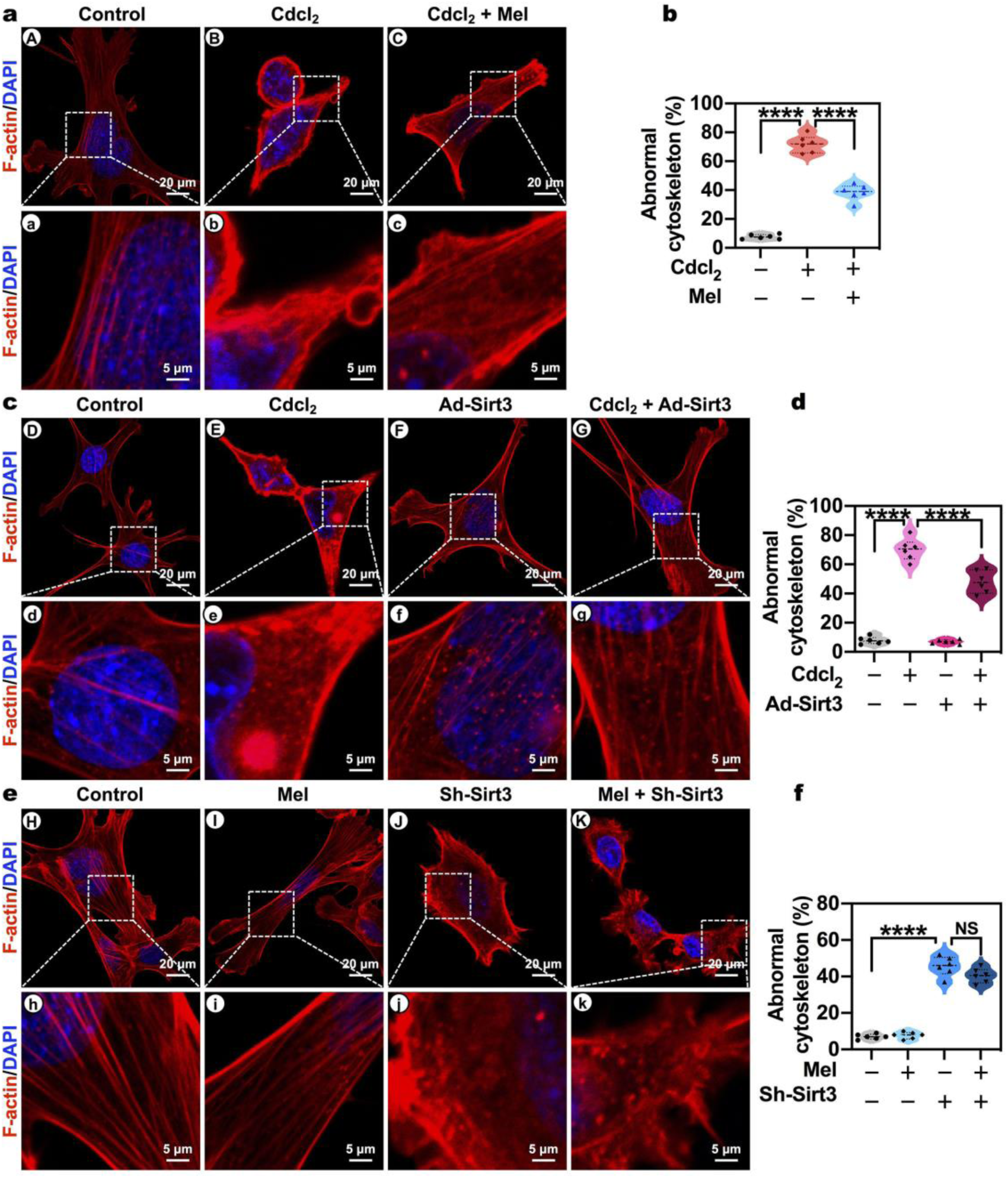
Overexpression and knockdown of Sirt3 protects and impairs the F-actin-containing cytoskeleton assembly in Mel-mediated TM4 mouse Sertoli cells protection. **a, c, e** Representative confocal microscope imaging of F-actin (a marker of cytoskeleton) in TM4 cells by the immunofluorescent analysis of phalloidin. Areas (A-K) in the upper panels (Scale bar, 20 μm) have been shown for further details (a-k) in the low panels (Scale bar, 5 μm). (A-C) Mel protects the cytoskeleton assembly in TM4 cells; (D-G) Sirt3 overexpression presents well-organized F-actin structure, and salvages the F-actin disruption induced by Cd; (H-K) the knockdown of Sirt3 apparently disturbs the F-actin network, which fails to be rescued by Mel. **b, d, f** The percentages of TM4 cells with abnormal F-actin-containing cytoskeleton per 200 cells. \**p* < 0.05; \*\**p* < 0.01; \*\*\**p* < 0.001; \*\*\*\**p* < 0.0001.

Overall, these results indicated that **SIRT3, in Sertoli cells, regulated SOD2 deacetylation and negative cytoskeleton protein PDLIM1 to orchestrate F-actin-containing cytoskeleton assembly, which accounted for basal ES (Sertoli cell-Sertoli cell interface) and apical ES (Sertoli cell-spermatid interface), even microtubule-based manchette in elongating spermatids.**

## Discussion

Mammalian spermatogenesis requires highly-coordinated Sertoli cell and Leydig cell for germ cell differentiation through mitosis, meiosis, and spermiogenesis (**Fig. 8**). The absence of this homeostasis impairs sperm count and motility, eliciting male infertility (Garolla et al., 2013; Wang et al., 2019). Mammalian spermatogenesis requires highly-coordinated Sertoli cells and Leydig cells for germ cells differentiation through mitosis, meiosis, and spermiogenesis. Here, we elucidate that SIRT3 exerts a dual role in testis: regulating cholesterol metabolism and cytoskeleton assembly during spermatogenesis. Mechanistically, in Leydig cells, phosphorylation of NF-κB p65^Ser536^ stimulates the nuclear translocation of NF-κB molecules (subunits p50, p65, RelB), which bind to the promoter of Sirt3; particularly, p50 represses Sirt3 transcription by simultaneously binding to TFBS1 and TFBS2 in Sirt3 promoter; then, inhibited SIRT3 disturbs cholesterol metabolism markers ABCA1, StAR and P450scc deacetylation (**Fig. 8(II)**). Whereas, in Sertoli cells, SIRT3 regulates SOD2 deacetylation and negative cytoskeleton protein PDLIM1 to orchestrate F-actin-containing cytoskeleton assembly, including microtubule, basal ES (Sertoli cell-Sertoli cell interface) in blood-testis barrier (BTB) via JAM-A, Occluding, N-Cadherin, β-catenin (**Fig. 8(III)**). SIRT3 mediates cytoskeleton assembly of elongating spermatids, including apical ES (Sertoli cell-spermatid interface), even microtubule-based manchette during spemiogenesis (**Fig. 8(I)**). To sum up, NF-κB-repressed SIRT3 acts directly on cholesterol metabolism of Leydig cells and cytoskeleton assembly of Sertoli cells via P450scc/SOD2 deacetylation to regulate sperm differentiation, influencing spermatogenesis, even male fertility.

**Fig. 8.**
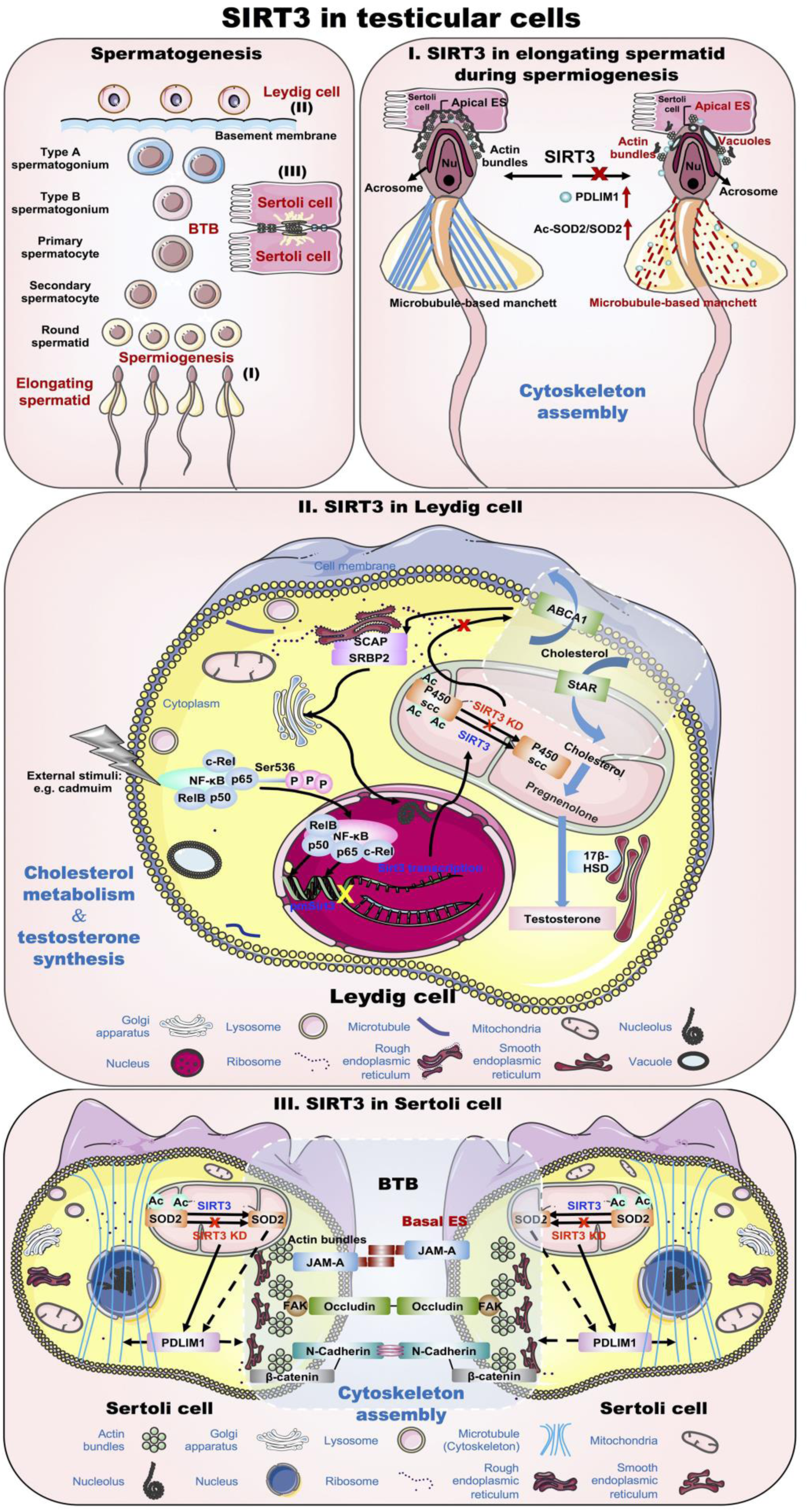
Summary diagram for the role of SIRT3 in testicular cells. Mammalian spermatogenesis requires highly-coordinated Sertoli cells and Leydig cells for germ cells differentiation through mitosis, meiosis, and spermiogenesis. Here, we elucidate that SIRT3 exerts a dual role in testis: regulating cholesterol metabolism and cytoskeleton assembly during spermatogenesis. Mechanistically, in Leydig cells, phosphorylation of NF-κB p65^Ser536^ stimulates the nuclear translocation of NF-κB molecules (subunits p50, p65, RelB), which bind to the promoter of Sirt3; particularly, p50 represses Sirt3 transcription by simultaneously binding to TFBS1 and TFBS2 in Sirt3 promoter; then, inhibited SIRT3 disturbs cholesterol metabolism markers ABCA1, StAR and P450scc deacetylation **(II)**. Whereas, in Sertoli cells, SIRT3 regulates SOD2 deacetylation and negative cytoskeleton protein PDLIM1 to orchestrate F-actin-containing cytoskeleton assembly, including microtubule, basal ES (Sertoli cell-Sertoli cell interface) in blood-testis barrier (BTB) via JAM-A, Occluding, N-Cadherin, β-catenin **(III)**. SIRT3 mediates cytoskeleton assembly of elongating spermatids, including apical ES (Sertoli cell-spermatid interface), even microtubule-based manchette during spemiogenesis **(I).** To sum up, NF-κB-repressed SIRT3 acts directly on cholesterol metabolism of Leydig cells and cytoskeleton assembly of Sertoli cells via P450scc/SOD2 deacetylation to regulate sperm differentiation, influencing spermatogenesis, even male fertility.

Cholesterol metabolism is prerequisite for testosterone biosynthesis in testicular Leydig cells (Gao et al., 2018). In vivo study, Mel, as SIRT3 activator (Zhai et al., 2017; Zhang et al., 2017), salvaged male reproductive function injury—sperm count and motility reduction, testosterone reduction and cholesterol augment of serum in Cd-induced subfertility rats model (**Fig. 1a-f**). Cholesterol and testosterone level mirror a dynamic balance of cholesterol metabolism: biosynthesis, uptake, export and esterification (Luo et al., 2020). Our observations about cholesterol metabolism markers that Cd-induced inhibition of Abca1 (cholesterol efflux marker), Scap/Srebf2 (cholesterol metabolism regulators) and Hsd17b3 (testosterone biosynthesis marker), and stimulation of Star (mitochondrial cholesterol influx transporter) were ameliorated by Mel in testis, corresponded to the changes in the level of cholesterol and testosterone. The results suggest that SIRT3 activator Mel participates in cholesterol metabolism, affecting testosterone production in testis.

Testosterone is essential for several crucial plots during spermatogenesis: BTB (basal ES/Sertoli-Sertoli adhesion), meiosis, apical ES (Sertoli-spermatid adhesion), and sperm release (Smith and Walker, 2014). Our data show that SIRT3 activator Mel rescued the structure disorganization of Sertoli cell, Leydig cell, and BTB induced by Cd, which accounts for the above male reproductive function injury. Apart from basal ES in BTB, cytoskeleton such as apical ES, microtubule and axoneme, are responsible for sperm head reshaping and flagella formation during spermiogenesis, promoting sperm motility acquirement (Pleuger et al., 2020). To identify whether Mel protects cytoskeleton assembly, facilitating sperm motility, we detected the dynamic process of spermiogenesis by TEM. Despite the normal axoneme, Cd undermined apical ES and microtubule-based manchette in elongating spermatids, which were alleviated by Mel. Parallelly, Mel rescued F-actin-containing cytoskeleton disruption induced by Cd. Molecularly, Mel-mediated cytoskeleton assembly is concomitant with the changes in BTB markers (Occludin, JAM-A, N-cadherin) (Cheng and Mruk, 2012), apical ES–BTB axis regulators (FAK, p-FAK-Tyr407) (Cheng and Mruk, 2012), and cytoskeleton negative regulators (PDLIM1) (Shang et al., 2016). Taken together, these results echo our hypothesis that Mel engages in cytoskeleton assembly, including basal ES, apical ES, and microtubule-based manchette during spermatogenesis.

Upregulation of SIRT3 following Mel has been demonstrated as a protective factor in a diversity of tissues and cells, such as myocardium (Zhai et al., 2017), hepar (Song et al., 2017), cerebrum (Liu et al., 2019), which is accordant with our data that Mel indeed stimulated SIRT3, expediting SOD2 deacetylation by directly interacting with SOD2 in testis. SIRT3-SOD2 axis regulates the scavenge of ROS (Kim et al., 2017), while the balance of ROS is required for actin cytoskeletal dynamics (Gourlay and Ayscough, 2005; Hunter et al., 2018). These data hint that Mel may regulate SOD2 deacetylation and cytoskeleton negative regulator PDLIM1 by SIRT3 in testicular cytoskeleton assembly.

Except for SOD2, Mel also regulated P450scc deacetylation by the interaction of SIRT3 and P450scc in testis. P450scc, as a catalyst converting cholesterol to pregnenolone, is indispensable for testosterone biosynthesis (Jing et al., 2020). Besides, the data showing that Mel reversed NF-κB p65^Ser536^ phosphorylation in Cd-treated rat testis, is consistent with the work by Liu et al., who manifested that Mel blocked NF-κB p65 signal in mice adipose tissue (Liu et al., 2017b). Actually, the inhibition of NF-κB p65 regulates testosterone production by Nur77 and SF-1 (Hong et al., 2004). Therefore, P450scc acetylation and NF-κB p65^Ser536^ phosphorylation are responsible for cholesterol metabolism and testosterone biosynthesis. Plus, Winnik et al. delineated that Sirt3 knockout mice failed to address a high-cholesterol diet, and resulted in endothelial cell dysfunction (Winnik et al., 2016), displaying that SIRT3 was associated with cholesterol metabolism in endothelial cells. These data and ours imply Mel may regulate NF-κB p65^Ser536^ phosphorylation and P450scc deacetylation by SIRT3 in testicular cholesterol metabolism.

In vitro study, to confirm the role of SIRT3 in cholesterol metabolism in Leydig cells, we generated Sirt3 overexpression and knockdown models in TM3 mouse Leydig cells (TM3 cells). Like Mel, Sirt3 overexpression in TM3 cells rescued cell apoptosis and cholesterol metabolism disruption induced by Cd; however, the knockdown of Sirt3 induced cholesterol metabolism disruption similar to Cd, which failed to be rescued by Mel, revealing that Mel-mediated cholesterol metabolism was dependent on Sirt3 in TM3 cells.

SIRT3 dominated P450scc deacetylation in TM3 cells, which is consistent with in vivo study. Strikingly, whether Sirt3 overexpression or Sirt3 knockdown conducted no influence on NF-κB p65^Ser536^ phosphorylation, despite Mel indeed reversed Cd-stimulated NF-κB p65^Ser536^ phosphorylation, suggesting that Mel regulated NF-κB p65^Ser536^ phosphorylation, which was not dominated by Sirt3 in TM3 cells. Since Sirt3 was not the upstream regulator of NF-κB, what’s the relationship between NF-κB and Sirt3? Our data displaying that NF-κB p65^Ser536^ phosphorylation stimulated nuclear translocation of NF-κB subunits p50, p65, and RelB, is supported by three studies pinpointing that NF-κB can translocate to the nucleus through Ser^536^ phosphorylation (Yu et al., 2020). Using bioinformatics analysis, we predicted that NF-κB could be the upstream transcription factor of Sirt3. Our ChIP-qPCR assay demonstrated the combination between Sirt3 promoter and NF-κB subunits (p50, p65, RelB), especially p50. A dual-luciferase reporter assay identified that p50 repressed Sirt3 transcription by binding to TFBS1 and TFBS2 in Sirt3 promoter (**Fig 5**). In contrast, Liu et al. deemed that NF-κB activated SIRT3 transcription in irradiated tumor cells (Liu et al., 2015). This paradox may be caused by the different types of cells. Taken together, Sirt3 transcriptional repression by NF-κB operates cholesterol metabolism via P450scc deacetylation in Leydig cells.

When it comes to cytoskeleton assembly, both basal ES (Sertoli-Sertoli cell interface) and apical ES (Sertoli-elongating spermatid interface) are associated with Sertoli cell. We investigate the role of SIRT3 in cytoskeleton assembly by Sirt3 overexpression and knockdown models in TM4 mouse Sertoli cells (TM4 cells). Similar to Mel, Sirt3 overexpression rescued F-actin-containing cytoskeleton assembly disruption and cell apoptosis in Cd-treated TM4 cells; however, the knockdown of Sirt3 elicited cytoskeleton assembly disruption, which failed to be rescued by Mel, indicating that Mel-mediated cytoskeleton assembly protection was dependent on Sirt3 in TM4 cells (**Fig 6, 7**). Molecularly, SIRT3 dominates SOD2 deacetylation by directly interacting with SOD2, and Mel-mediated SOD2 deacetylation depends on SIRT3, in response to in vivo study. In terms of PDLIM1, both Mel and the overexpression of Sirt3 extinguished the augment of PDLIM1 protein level induced by Cd; the knockdown of Sirt3 evoked PDLIM1 accumulation, impairing cytoskeleton assembly, which didn’t be reversed by Mel. Additionally, the overexpression of Pdlim1 has been demonstrated to disturb cytoskeleton assembly in Sertoli cells (Liu et al., 2016). SIRT3-SOD2-mediated ROS clearance is indispensable for cytoskeleton assembly (Gourlay and Ayscough, 2005; Hunter et al., 2018). These data and ours support the assumption that SIRT3 controls cytoskeleton assembly by PDLIM1 and SOD2 deacetylation in Sertoli cells.

Although the knockdown of Sirt3 by shRNA exerts an effective interference in TM3 (mouse Leydig) cells and TM4 (mouse Sertoli) cells, more high-efficiency methods such as the conditional knockout (cKO) of Sirt3 in Leydig cells and Sertoli cells should be involved in future study. Our models in TM4 cells can partially respond to the ectoplasmic specialization (ES) assembly including apical/basal ES, but the phenotype about ES in Sirt3 cKO mice will be required to confirm or refute this hypothesis. Meanwhile, the accurate mechanism by which SIRT3 regulates microtubule-based manchette remains to be elucidated in elongating spermatids. In addition, the direct modes of actions between SIRT3 and cholesterol metabolism markers (Abca1, Scap/Srebf2, Hsd17b3, and Star) deserve intensive investigation, which may offer a novel orientation for exploring cholesterol metabolism during spermatogenesis.

Overall (**Fig. 8**), the most striking finding of this study is that SIRT3 exerts a dual role in testis: Sirt3 transcriptional repression by NF-κB orchestrates cholesterol metabolism via P450scc deacetylation in Leydig cells; whereas, in Sertoli cells, SIRT3 regulates cytoskeleton (basal ES, microtubule) assembly via PDLIM1 with SOD2 deacetylation; the regulation of Leydig cells and Sertoli cells may account for apical ES and microtubule-based manchette assembly in elongating spermatids, influencing sperm differentiation during spermiogenesis. Distinct molecular mechanisms establish a novel signaling network, highlighting the complex and multi-faceted action of SIRT3 as robust deacetylase in testicular cells. Understanding the role of SIRT3 in regulating cholesterol metabolism and cytoskeleton assembly during spermatogenesis may allow for the development of additional rational combination therapies using SIRT3 activators such as Mel to improve therapies for male subfertility, even infertility.

## Methods

### Chemicals and reagents

Cdcl_2_ was from Sinopharm (CFSR-10005416, Shanghai, China). Melatonin (Mel) was from MedChemExpress (Cat#HY-B0075, USA). Testosterone ELISA kit was from Goybio (Cat#GOY-088B, Shanghai, China). Total and free cholesterol assay kits were from Applygen Technologies (Cat#E1015-50, Cat#E1016-50, Beijing, China). TRIzol reagents were from Invitrogen. PrimeScript RT-PCR and SYBR Premix ExTaq Kits were from Takara (Cat#2641A, Cat#RR420, Japan). BCA protein assay and ECL kits were from Beyotime Institute of Biotechnology (Shanghai, China). N-Cadherin, β-Catenin, Occludin, JAM-A, β-Actin, SOD2, SOD2(acetyl K68), PDLIM1, NUDC, GAPDH antibodies were from Abcam (Cat#ab18203, Cat#ab68183, Cat#ab167161, Cat#ab125886, Cat#ab8226, Cat#ab13534, Cat#ab137037, Cat#ab129015, Cat#ab109318, Cat#ab37168, Cambridge, UK); FAK, p-FAK-Tyr407 antibodies were from Bioss (Cat#bs-1340R, Cat#bs-3164R, MA, USA); SIRT3, P450scc, Acetylated-Lysine, NF-κB p50, NF-κB p65, Phospho-NF-κB p65^Ser536^, RelB, c-Rel, normal IgG antibodies from Cell Signaling Technology (Cat#5490S, Cat#14217S, Cat#9441S, Cat#3035S, Cat#8242S, Cat#3033S, Cat#10544S, Cat#4727T, Cat#2729S, MA, USA). Co-IP and ChIP kits were from Thermo Fisher Scientific (Cat#26140, Cat#26156, MA, USA). CCK-8 Kit was from Dojindo (Kumamoto, Japan). An Annexin V-FITC and PI Detection Kit was from BD Biosciences (New Jersey, USA). DMEM was from HyClone (Logan, Utah, USA). Collagenase and FBS were from Gibco (Australia). FITC Phalloidin FITC and TRITC Phalloidin rhodamine were from Yeasen (Cat#40735ES75, Cat#40734ES75, Shanghai, China).

## Experimental design

### Animal model

2-mo-old adult male SD rats (230 ± 30 g) were purchased from Tongji Medical College Animal Center. Animals were adapted for 7d to the new environment, and fed ad libitum. The conditions were maintained as follows: a 12-h light/dark phase, temperature (22–26℃) and humidity (50 ± 5%). Thirty-six rats were randomly divided into 3 groups. Restricted randomization was not applied. According to our previous subfertility/infertility model (Wang et al., 2020), all rats except controls were intraperitoneally injected with Cdcl_2_ (0.8 mg/ kg) for consecutive 7 d. Some rats were intraperitoneally injected with Mel (2 mg/kg) (SIRT3 activator (Zhai et al., 2017),(Song et al., 2017),(Liu et al., 2019)) at 2 h before Cd treatment. Given that both CdCl_2_ and Mel were dissolved in 0.9% NaCl, the control group was treated with 0.9% NaCl. All the animal experiments were permitted by the IACUC of Tongji Medical College, Huazhong University of Science and Technology (**Supplementary file**) (Permit IACUC Number: 2061), and were implemented ethically as the Guide for the Care and Use of Laboratory Animal guidelines.

### Cell models

TM3 mouse Leydig cells and TM4 mouse Sertoli cells were from the Institute of Reproductive Health, Tongji Medical College. After tested for mycoplasma contamination, TM3 and TM4 cells were respectively cultured in DMEM, which was supplemented with 10% FBS, at 37°C in a humidified atmosphere with 5% CO2/95% air. According to our previous study (Wang et al., 2020), IC50 of Cd for TM3 cells—8.725 μg/ml and IC50 of Cd for TM4 cells—12 μg/ml were exploited in subsequent cell models. Considering that 10 μM Cdcl_2_ could be rescued by 1 μM Mel in HepG2 cells, we determined 1.1 μg/ml Mel to protect against 8.725 μg/ml Cdcl_2_ in TM3 cells and 1.5 μg/ml Mel to protect against 12 μg/ml Cdcl_2_ in TM4 cells by unit conversion (**Supplementary Table 1**).

**Supplementary Table 1.**
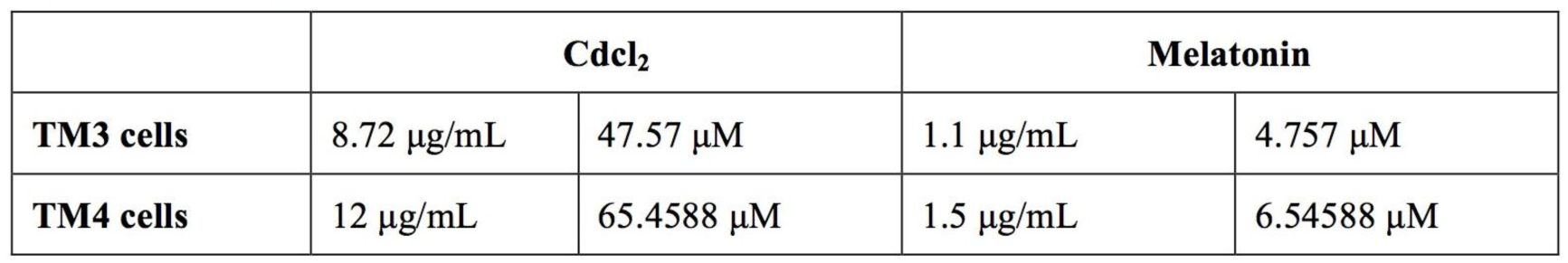
Unit conversion. (https://www.promega.com/resources/tools/biomath/molarity-calculator/)

**Supplementary Table 2.**
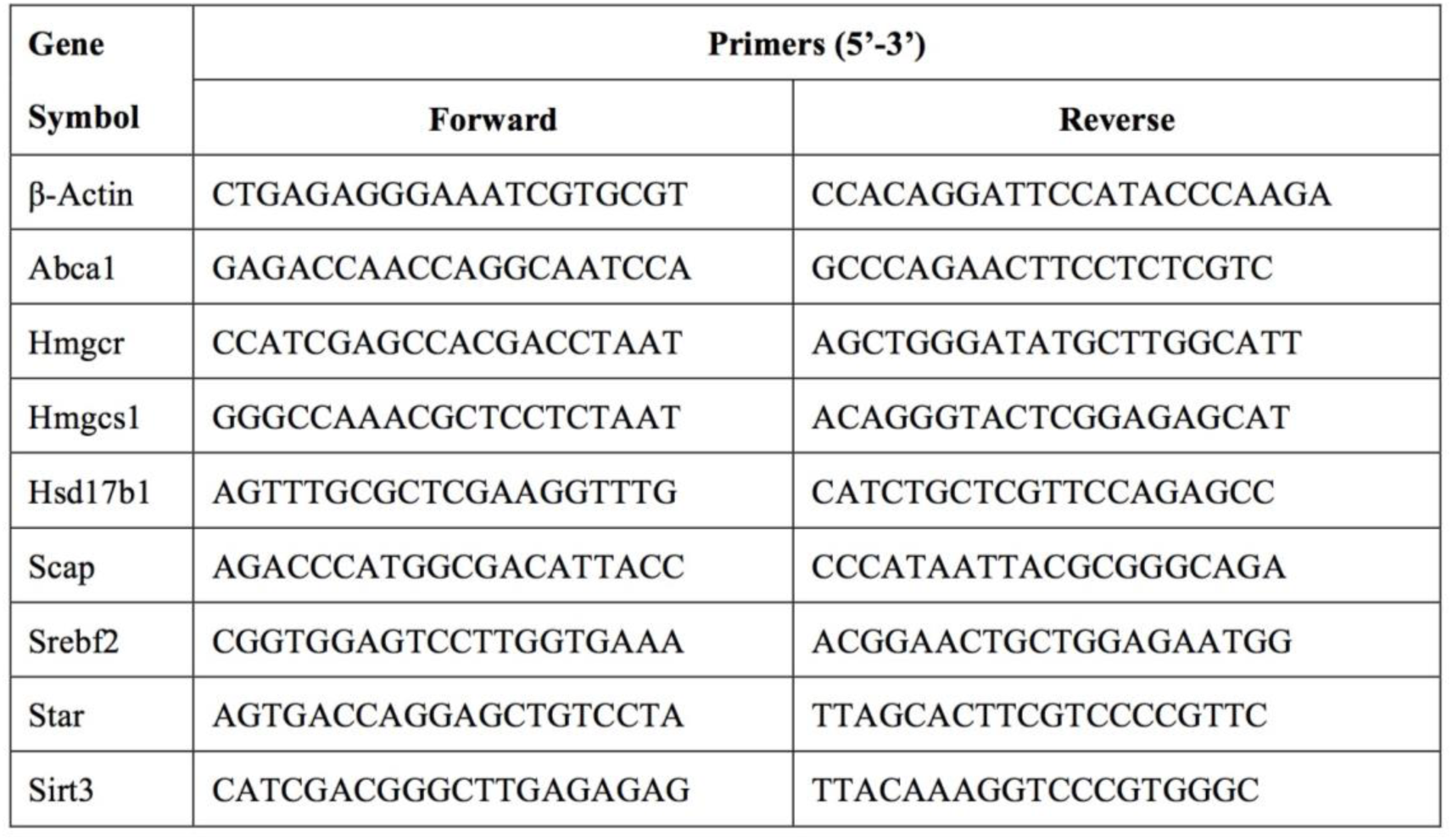
Sequences of the primers for quantitative real time PCR of mRNA levels.

To explore if Mel alleviated the Cd-induced TM3 cells injury, TM3 cells were randomly divided into 3 groups in model 1: control, Cdcl_2_ (8.725 μg/ml), Cdcl_2_ (8.725 μg/ml) with Mel (1.1 μg/ml). To identify if SIRT3 alleviated the Cd-induced TM3 cells injury, TM3 cells were randomly divided into 4 groups in model 2: control, Cdcl_2_ (8.725 μg/ml), Sirt3 overexpression (Ad-Sirt3), Cdcl_2_ (8.725 μg/ml) with Sirt3 overexpression (Ad-Sirt3). To examine if Mel-mediated cell protection was dependent on SIRT3, TM3 cells were randomly divided into 4 groups in model 3: control, Mel (1.1 μg/ml), Sirt3 knockdown (sh-Sirt3), Mel (1.1 μg/ml) with Sirt3 knockdown (sh-Sirt3).

In parallel, to investigate the role of SIRT3 in TM4 cells, TM4 cells were randomly divided into 3 models—TM4-model 1: control, Cdcl_2_ (12 μg/ml), Cdcl_2_ (12 μg/ml) with Mel (1.5 μg/ml); TM4-model 2: control, Cdcl_2_ (12 μg/ml), Sirt3 overexpression (Ad-Sirt3), Cdcl_2_ (12 μg/ml) with Sirt3 overexpression (Ad-Sirt3); TM4-model 3: control, Mel (1.5 μg/ml), Sirt3 knockdown (sh-Sirt3), Mel (1.5 μg/ml) with Sirt3 knockdown (sh-Sirt3).

24 h post Ad-Sirt3 or sh-Sirt3 transfection, some cells were exposed to Cd for 24 h. Some cells were pretreated with Mel for 2 h prior to Cd treatment. Researchers and statistical analysts were blind to the allocation of groups.

### Sirt3 overexpression and knockdown by adenovirus vectors

Mouse Sirt3 (NM_001177804.1) overexpression adenovirus were synthesized by a pAdM-FH-GFP vector (Vigene Biosciences) (**Supplementary Fig.1a**). The construct was confirmed by DNA sequencing. Sirt3 knockdown adenovirus was designed according to four sequences of shRNA as follows: (5′ → 3′ orientation) Sirt3-shRNA1: GGCTCTATA CACAGAACATCGTTCAAGAGACGATGTTCTGTGTATAGAGCCTTTTTT; Sirt3-shRNA2: GGCAATCTAGCATGTTGATCGTTCAAGAGACGATCAACATG CTAGATTGCCTTTTTT; Sirt3-shRNA3: AGACAGCTCCAACACGTTTACTTCA AGAGAGTAAACGTGTTGGAGCTGTCTTTTTTT; Sirt3-shRNA4: GCGTTGTGA AACCCGACATTGTTCAAGAGACAATGTCGGGTTTCACAACGCTTTTTT, which were constructed in a pAdM-4in1-shRNA-GFP vector (Vigene Biosciences) (**Supplementary Fig.1b**). Sirt3 overexpression and knockdown adenoviruses were respectively transfected into TM3 cells or TM4 cells with ADV-HR (FH880805) (Vigene Biosciences) for the Sirt3 overexpression or knockdown assay. The efficiencies of Sirt3 overexpression and knockdown were examined in TM3 cells and TM4 cells (**supplementary Fig. 1c, 1d**).

### Plasmid constructions

Mouse Nfkb1 (NM_008689.2) coding sequence was synthesized by gene synthesis (General Bio), and cloned into the pcDNA3.1 eukaryotic expression vector (**Supplementary Fig. 5a**). Sirt3-WT promoter and the mutant promoters Sirt3-MUT1 (deletion of TFBS1), Sirt3-MUT2 (deletion of TFBS2), Sirt3-MUT3 (simultaneous deletion of TFBS1 and TFBS2) (**Fig. 5f**) were constructed by PCR-based amplification and cloned into pGL3-Promoter vector (**Supplementary Fig. 5b**). Sequences of the primers for amplification of promoters are shown in **Supplementary Table 3**. All constructs were confirmed by DNA sequencing (**Available from authors**). NF-κB1 protein coding capacity in HEK293T cells was tested by western blot assay (**Supplementary Fig. 5d**).

**Supplementary Table 3.**
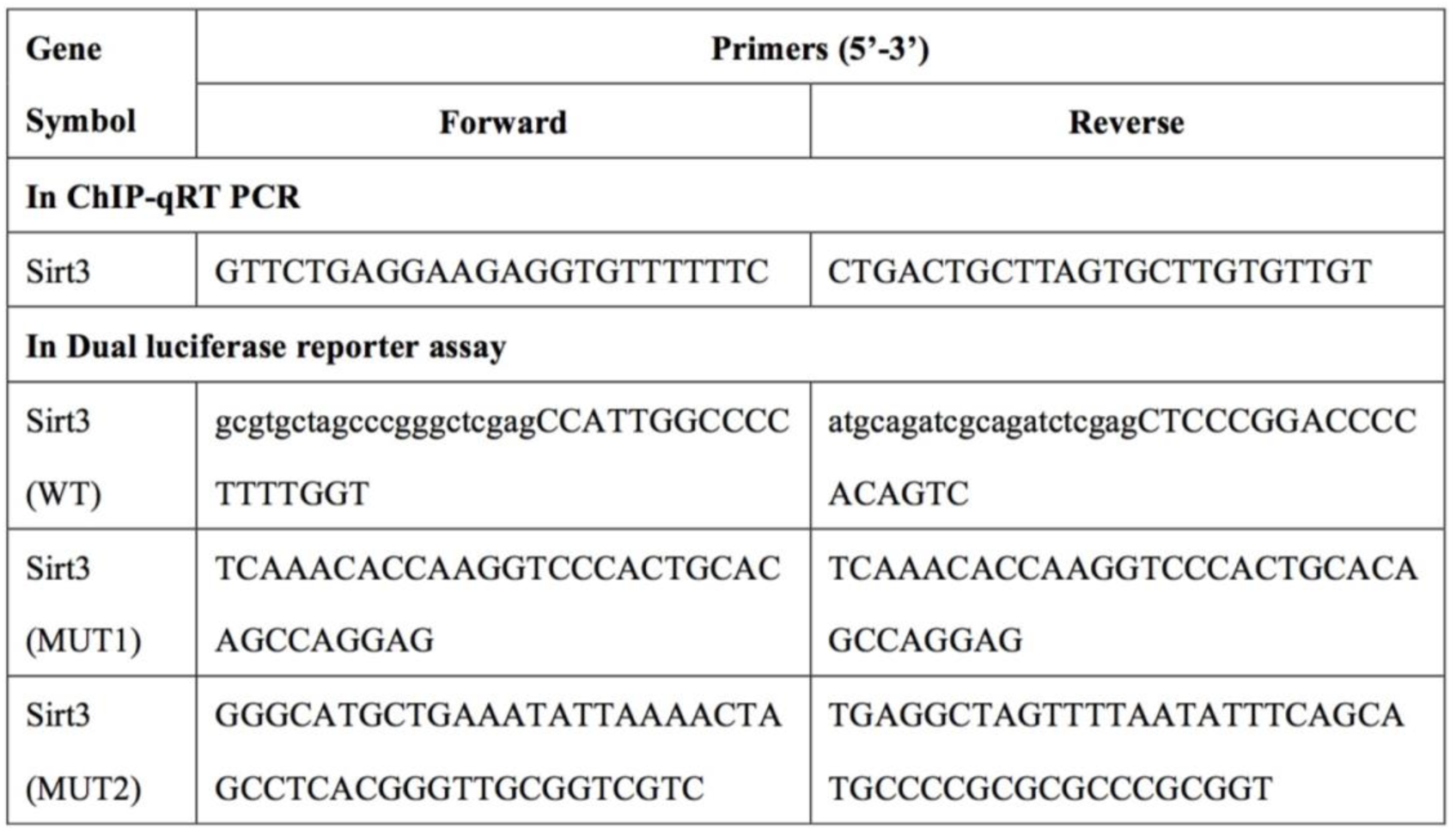
Sequences of the primers for amplification of promoter.

**Supplementary Fig. 5.**
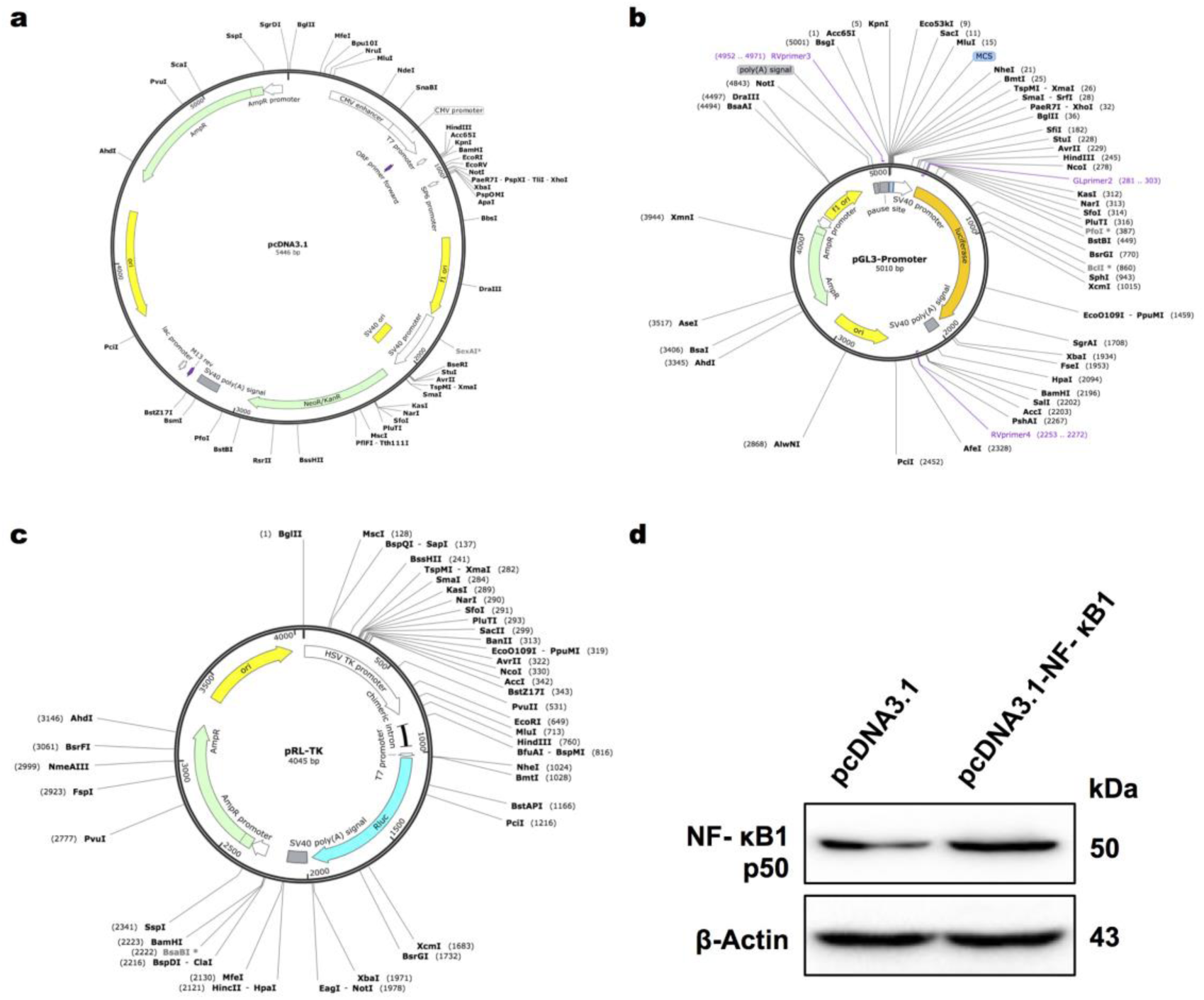
Plasmid constructions for dual luciferase reporter assay. **a** pcDNA3.1 eukaryotic expression vector for overexpression of NF-κB1 in HEK293T cells. **b** pGL3-Promoter vector for insertion of Sirt3 wild-type or mutant promoter in HEK293T cells. **c** pRL-TK vector as an internal control for transfection. **d** NF-κB1 protein coding capacity in HEK293T cells by western blot assay.

### Dual-luciferase reporter assay

By Lipofectamine® 3000 transfection reagent (Invitrogen), HEK293T cells (from Tongji Medical College) were transiently transfected with the NF-κB p50 expression vector (pcDNA3.1-NF-κB), or empty control vector as well as WT or TFBS1/TFBS2 mutant Sirt3 promoter (MUT1/MUT2/MUT3) luciferase reporter vector (pGL3-Promoter) and Renilla luciferase-reporter vector (pRL-TK, as an internal control for transfection) (**Supplementary Fig. 5c**). Luciferase activities were determined after 24-h incubation with the Dual-luciferase reporter assay system (Promega). The firefly luciferase activity was normalized to renilla luciferase activity. The relative luciferase activity was analyzed as a function of NF-κB p50-dependent Sirt3 transcription.

### Chromatin immunoprecipitation (ChIP) assay

The ChIP assay was implemented by a ChIP assay kit according to the manufacturer’s instructions (Thermo Fisher Scientific). 1% formaldehyde was utilized to cross-link the histones and genomic DNA in TM3 mouse Leydig cells. Cell lysates were prepared, and chromosomal DNA was sonicated to obtain average sizes between 200 and 1000 bp. The chromatin was incubated and precipitated with antibodies against p50, p65, RelB or normal rabbit IgG (CST) as controls at 4 °C overnight. DNA fragment that contained the NF-kB binding site of the Sirt3 promoter was amplified. The amplified products were analyzed by quantitative real-time PCR.

### Sperm parameter analysis

According to “WHO laboratory manual for the examination and processing of human semen (Fifth edition, 2010)”, the sperm count was calculated by multiplying the semen volume and concentration, which was measured via an improved Neubauer hemocytometer. Sperm motility was analyzed through the progressive sperm count per 200 sperm under a microscope.

### Cd level in the testis

Cd level in testis was detected by graphite furnace atomic absorption spectrometry (GFAAS) as the previous method (Wang et al., 2020).

### Testosterone and cholesterol levels

Testosterone levels in rat serum and cultured TM3 cells supernatants were qualified by testosterone ELISA kit following the manufacturer’s instructions (Goybio). Total cholesterol (TC) levels in testis and TM3 cells were measured by total cholesterol assay kit; free cholesterol (FC) levels were determined by free cholesterol assay kit according to the manufacturer’s instructions (Applygen Technologies).

### Quantitative real-time PCR assay

The mRNA expression levels of Abca1, Hmgcr, Hmgcs1, Hsd17b1, Scap, Srebf2, Star in testis and TM3 cells, and Sirt3 in TM3/TM4 cells were analyzed by quantitative real-time PCR assay. SYBR RT-PCR kit (Takara) and LightCycler (Roche) were exploited according to the standard procedures. Data were normalized to β-actin expression. Sequences of the primers for quantitative RT-PCR are in **supplementary Table 2**.

### Histopathological analyses

Testes were fixed in Bouin’s fixative for 24 h, and embedded in paraffin, and sectioned. Testes sections were stained with hematoxylin and eosin (H&E) and observed under a light microscope for structure.

### Transmission electron microscopy (TEM)

Testes were fixed with 2.5% glutaraldehyde for 2 h at 4 °C, postfixed in 1% osmium tetroxide, and embedded in Epon 812 as the previous method (Wang et al., 2020); the ultrastructure of Sertoli cell, Leydig cell, BTB, and cytoskeleton during spermiogenesis in testis was investigated by TEM (JEM 1200-EX; Hitachi, Ltd, Tokyo, Japan) at 80 kV.

### F-actin examination by immunofluorescence and confocal microscopy

Fixed testis sections were stained with FITC Phalloidin FITC (Yeasen), which labeled F-actin. TM4 cells were washed and fixed with 4% paraformaldehyde, and then stained with TRITC Phalloidin rhodamine (Yeasen), which labeled F-actin. Under Nikon A1 laser scanning confocal microscope (Nikon America Inc., Melville, NY), F-actin structure was investigated. The percentages of spermatids or Sertoli cells with abnormal F-actin-containing cytoskeleton per 200 elongating spermatids or TM4 cells were calculated in 6 independent experiments for each group.

### Western blotting

The protein concentration was detected by a BCA protein assay kit. Proteins were denatured, separated on SDS-PAGE gels, and transferred to a PVDF membrane as the previous method (Wang et al., 2020). The membranes were incubated with antibodies according to the manufacturer’s instructions, including N-Cadherin, β-Catenin, Occludin, JAM-A, β-Actin, SOD2, SOD2(acetyl K68), PDLIM1, NUDC, GAPDH, FAK, p-FAK-Tyr407, SIRT3, P450scc, Acetylated-Lysine, NF-κB p50, NF-κB p65, Phospho-NF-κB p65^Ser536^, RelB, c-Rel. β-Actin and GAPDH proteins were used as a loading control for total and cytoplasmic proteins respectively. The membranes were incubated with corresponding secondary antibodies; the target proteins were detected with ECL; the density was analyzed by Image J.

### Co-immunoprecipitation (Co-IP) assay

co-IP was performed by the Thermo Scientific Pierce co-IP kit according to the manufacturer’s protocol (Thermo Fisher Scientific). For the detection of interaction between SOD2 and SIRT3, 60 μl of anti-SOD2 antibody, 60 μl of anti-SIRT3 antibody, and 30 μl of IgG antibody were first immobilized for 2 h using AminoLink Plus coupling resin, which was washed and then incubated with 200 μl (500 μg of proteins) of lysate. Next, the resin was precleaned with control agarose resin for 1 h and incubated. Using the elution buffer, the coupling resin was again washed and protein was eluted. For the detection of interaction between P450scc and SIRT3, the method was as above. Samples were analyzed by immunoblotting.

### CCK-8 assay

Cell viabilities of TM3 and TM4 cells were assessed by CCK-8 assay kit according to the manufacturer’s instructions as the previous method (Wang et al., 2020).

### Flow cytometry analysis

To analyze the effects of the indicated treatments on cell survival, we stained the cells with an Annexin V-FITC and PI Detection Kit and analyzed them by flow cytometry; flow cytometry data were assessed using BD FACSDiva Software v7.0 (Becton-Dickinson, USA) (Wang et al., 2020).

### Statistical analysis

The data were expressed as the mean ± S.D. Differences among two groups were analyzed by Student’s *t*-test, the Mann–Whitney *U*-test, or Generalized estimating equation (if non-normal distribution). Differences among multiple groups were analyzed by one-way analysis of variance (ANOVA). Multiple comparisons for subgroups were determined by Dunnett’s T3 tests. Statistical significance was considered as follows: NS, *p* > 0.05; *, &, #, *p* < 0.05; \*\**p* < 0.01; \*\*\**p* < 0.001; \*\*\*\**p* < 0.0001. The showed experiments were replicated 4-10 times.

## Data availability

All data generated or analyzed during this study are included in the manuscript and supporting files. Source data files have been provided for Figures 1-7 and supplementary files.

## Acknowledgements

The authors thank the Editors and Reviewers for their significant contributions during the revision period. This study was supported by National Key R&D Program of China (No. 2020YFA0803900), Hubei Science and Technology Plan (No. 2017ACB640), and Wuhan University Medical Development Plan (No. TFJC2018001).

## Author contributions

M.W, Y.Z.Z. and P.S. conceived the project and designed the experiments; M.W., L.Z., Y.X., X.F.W., L.C. performed the experiments; M.W, F.W., analyzed the data, and wrote the paper; M.W., and Y.X. edited the manuscript; Y.Z.Z. and P.S. supervised the study; Y.Z.Z. obtained fundings for the work.

---

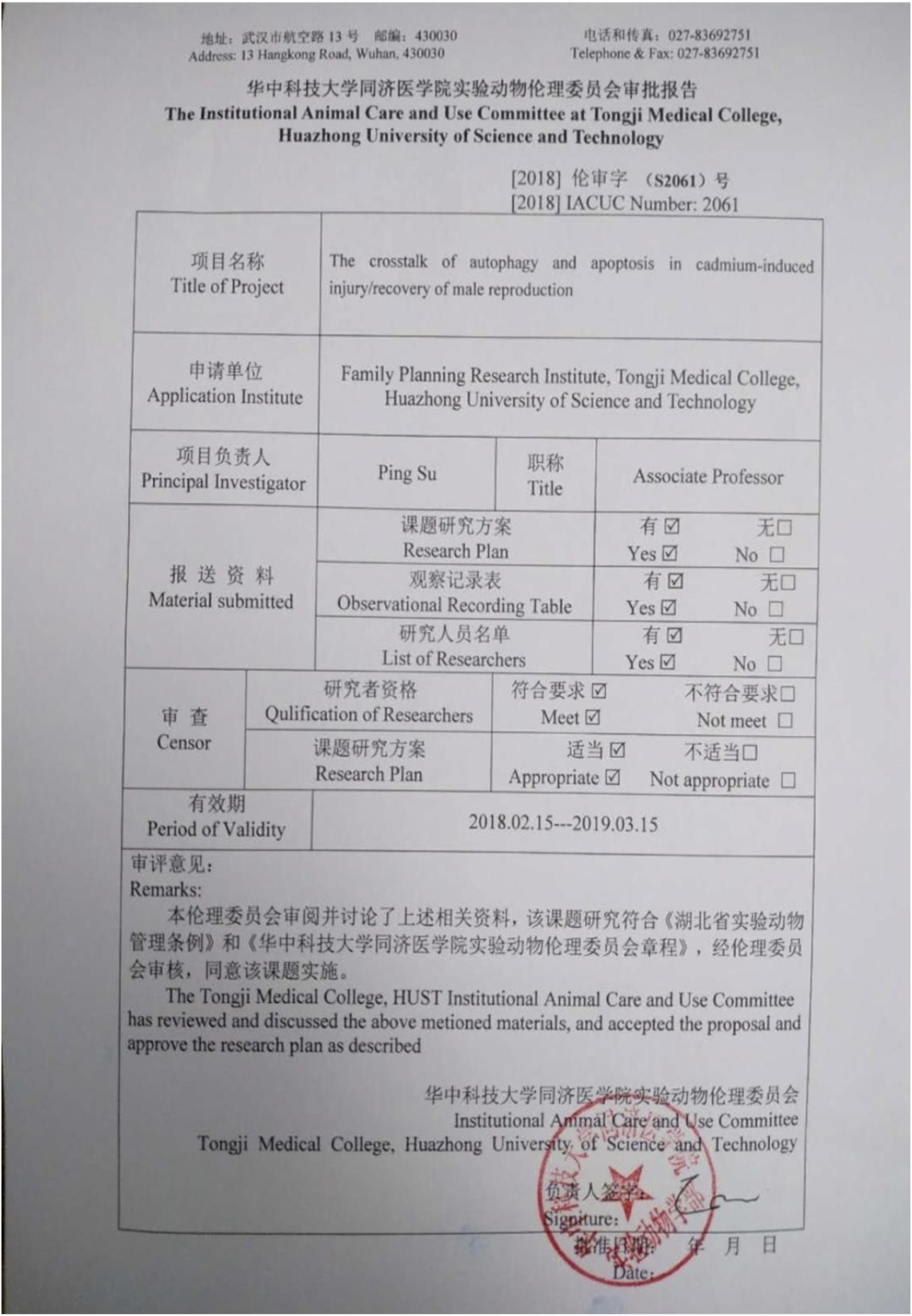

